# A thalamic hub integrates brainstem and stress signals to dynamically regulate REM sleep

**DOI:** 10.1101/2025.07.22.665932

**Authors:** Aletta Magyar, Péter Berki, Sándor Borbély, Boglárka Barsy, Anna Virág Bakacsi, Magor L Lőrincz, Danqian Liu, Ferenc Mátyás

**Author notes:** Corresponding author: Ferenc Mátyás.

## Abstract

Rapid eye movement (REM) sleep, a brain state critical for sleep quality and cortical cognition, is tightly controlled by brainstem circuits and highly sensitive to stress. Yet, how these systems interact to regulate REM sleep and its associated forebrain rhythms remains elusive. Here, we identify a thalamic hub that conveys medullary REM-promoting signals, and integrates stress inputs to regulate REM sleep. A subpopulation of paraventricular thalamic neurons that collaterally project to the cortex and nucleus accumbens (PVT_→NAc_) selectively responds to activation of medulla REM-promoting neurons, and bidirectionally modulates REM-associated theta oscillations in an activity-dependent manner. Their low-frequency activation promotes theta rhythms during REM sleep, while high-frequency activation suppresses them, mirroring the neuronal signatures of acute and chronic stress, respectively. Distinct patterns of PVT_→NAc_ neurons underlie the bidirectional stress modulation of REM sleep, partly by differentially engaging prefrontal microcircuits via their collateral projections. Together, our findings uncover a thalamic integrative hub that couples sleep-regulating and stress pathways to adaptively control REM sleep expression.

## Introduction

Neuronal oscillations of cortical networks are fundamental for regulating brain states and influencing cognitive functions. In particular, Rapid Eye Movement (REM) sleep has been linked to key cognitive processes such as memory formation and problem solving, in both human and mice^1–5^. Furthermore, disruption of REM sleep and the associated theta oscillation (4-8 Hz) has been implicated in various psychiatric disorders, including post-traumatic stress disorder, anxiety disorders, cognitive impairments in Parkinson’s disease as well as Alzheimer’s disease^1,6–9^. These findings suggest that cortical REM rhythm could serve as valuable biomarker for the development of such pathologies.

REM sleep has been shown to be regulated by brain-wide neuronal networks ^10–13^. Brainstem circuits, including the medullar GABAergic signals, have been identified as crucial executive hubs for generating REM sleep and cortical theta oscillation^13–16^. Recent evidence has highlighted that cortical regions, whose excitation–inhibition balance is altered during REM sleep, is directly involved in the regulation of REM sleep^17,18^. However, in the absence of direct cortical innervation, it remains unclear how medullar ascending signals are transmitted to forebrain circuits to modulate REM sleep theta oscillations, and consequently, regulate REM states.

The stress-sensitive nature of REM sleep implies the existence of a neural switching system capable of integrating sleep-, salience- and stress-related information from brainstem and midbrain structures^19,20^. Cortical functions related to sleep and cognition are strongly influenced by its thalamic inputs^21–23^, particularly those originating from the midline thalamic paraventricular nucleus (PVT)^24–26^. Neuronal activity of PVT plays a key role in encoding salience, arousal and homeostatic signals within the forebrain^25–29^, as well as in regulating behavioral responses to stress^30–34^. Notably, the PVT receives broad ascending inputs from the brainstem and sends extensive projections to the prefrontal cortex (PFC). This positions the PVT as a candidate link between medullary and cortical mechanisms of REM sleep that can integrate arousal and stress information.

Thus, in this work, by combining circuit tracing, optogenetic manipulation, and cell-type-specific electrophysiological recordings, we identified the engagement of the PVT in the regulation of REM sleep and the associated theta oscillation at the level of PFC operation. Our data show that a subpopulation within the PVT that send collateral projections to the PFC and nucleus accumbens (PVT_→NAc_) was selectively activated by direct REM-promoting VLM_GABA_ inputs, and display REM sleep-associated activation. The activity of PVT_→NAc_ cells not only regulates theta oscillations and the associated excitation/inhibition balance in the PFC during physiological sleep, but also recruits adaptive mechanisms in stress-dependent manner. Taken together, our findings uncover a brainstem–thalamus–cortex circuit through which the paraventricular thalamus dynamically regulates REM sleep and cortical theta oscillations, providing a framework for understanding how sleep and stress interact to shape higher-order brain functions.

## Results

### PVT transmits VLM_GABA_ signal to the PFC

We first examined whether activation of the REM-promoting ventrolateral GABAergic neurons (VLM_GABA_) could affect cortical theta oscillations, as it was earlier reported in drug-free condition^14^, by combining cell-type-specific optogenetics and electrophysiological recordings in the PFC. Specifically, VLM_GABA_ cells were virally transduced with Cre-dependent recombinant adeno-associated virus (AAV) expressing Channelrhodopsin 2 (ChR2) in vGAT-Cre mice (Fig. 1a). After 3-4 weeks of recovery, VLM_GABA_ neurons were optogenetically stimulated, which – indeed – evoked an increase in cortical theta oscillatory activity in all tested mice, measured in the 4-8 Hz frequency band (Fig. 1b and Extended Data Fig. 1, also see *Methods*).

**Fig. 1:**
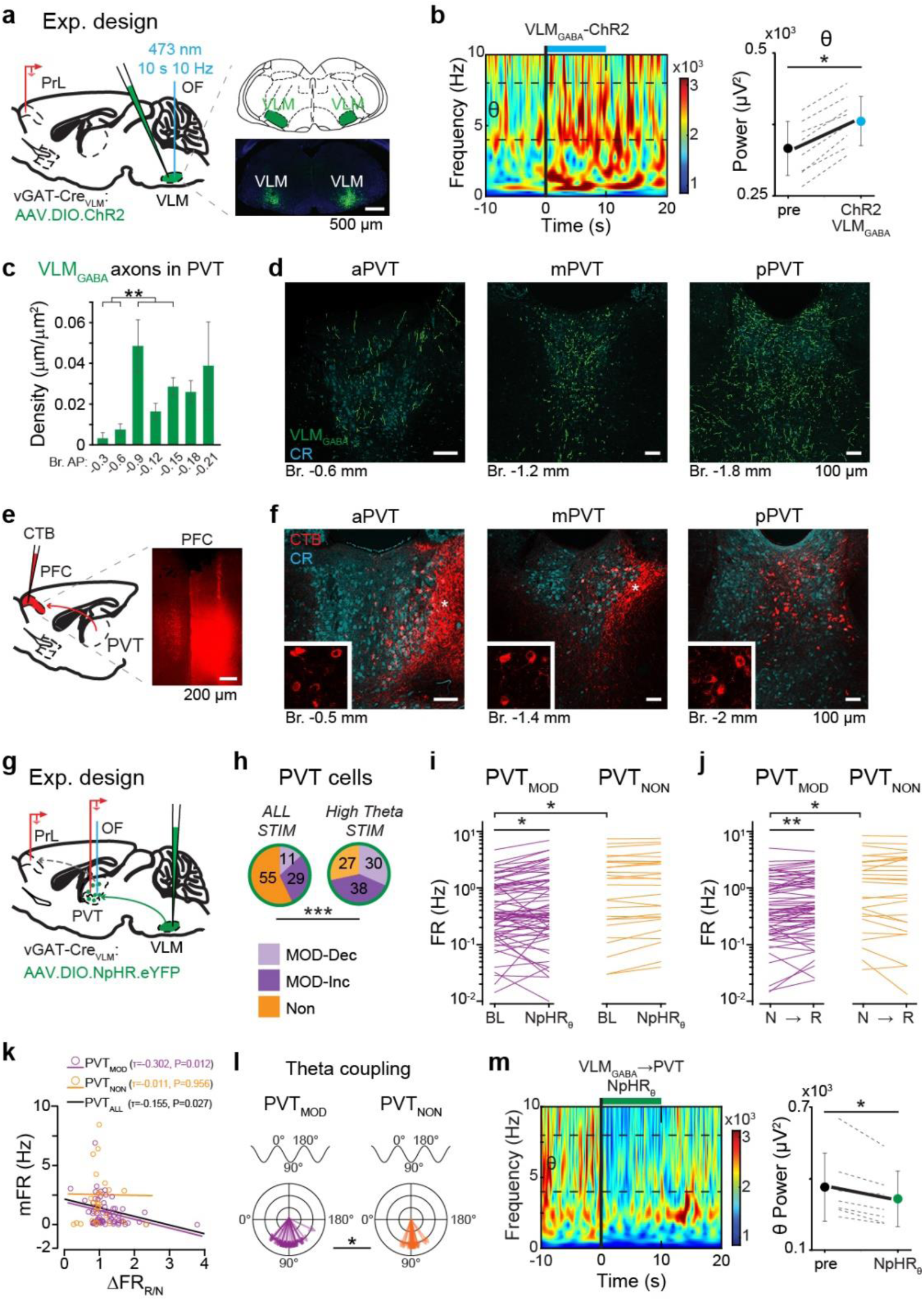
VLMGABA shape REM-related cortical theta oscillation via PVT. **a,** *Left,* experimental design for prefrontal cortical LFP recordings with optogenetic activation of VLM_GABA_ neurons. *Right*, injection sites targeting VLM_GABA_ cells with AAV-DIO-ChR2 in vGAT-Cre mice. **b,** *Left*, effect of VLM_GABA_ stimulation on the LFP spectrogram. *Right*, population data for theta (θ) band power changes induced by VLM_GABA_ stimulation (n = 8 animals; Wilcoxon matched-pairs test). **c,** Quantification of topographic distribution of VLM_GABA_-to-PVT inputs (n = 3 animals; Shapiro-Wilk W test with paired t-test). **d,** Representative confocal images of eYFP-labeled VLM_GABA_-to-PVT (green) at three anteroposterior levels. Cyan color shows the CR counterstaining indicating territory of PVT. **e,** Schematic (*left*) of and injection site (*right*) targeting the PFC with retrograde (CTB) tracer. **f,** Representative confocal images of CTB-labeled PFC-projecting PVT cells (red) at three anteroposterior levels corresponding to **d**. Cyan color shows the CR counterstaining indicating territory of PVT. Insets, high magnification images of CTB labeled cells. Asterisks indicate cortico-thalamic fibers. **g,** Experimental design for simultaneous PVT and PFC recordings with optogenetic silencing (NpHr3.0) of VLM_GABA_-to-PVT inputs. **h,** Pie charts showing state-dependent responsiveness of PVT cells (N = 95; n = 7 mice) to inhibition of VLM_GABA_-to-PVT inputs (Pearson’s χ2 test). *ALL STIM,* all trials; *High Theta STIM,* inhibition during high theta power (REM-like) periods. **i,** Baseline (BL) and evoked PVT activity by inhibition of VLM_GABA_-to-PVT inputs during high theta trials (NpHR_HT_) clustered by responsiveness to the inhibition [PVT_MOD_ (purple) vs. PVT_NON_ (orange)] [Wilcoxon matched-pairs test for BL vs. NpHR_HT_; Mann-Whitney U test for PVT_MOD_ (N=68 cells) vs. PVT_NON-MOD_ (N=27 cells)]. **j,** Firing rate changes upon NREM-like (N)-to-REM-like (R) state transition were observed only in PVT_MOD_ cells (Wilcoxon matched-pairs test for NREM vs. REM; Mann-Whitney U test for PVT_MOD_ vs. PVT_NON_). **k,** Negative correlation between mean firing rate (mFR) and high theta-induced firing rate changes (R/N) observed only in PVT_MOD_ cells (Kendall’s TAU Correlations). **l,** Vector diagrams represent distinct, phase-locked firing of PVT_MOD_ and PVT_NON_ cells during theta waves (Pearson’s χ2 test). **m,** *Left*, Effect of optogenetic silencing of VLM_GABA_-to-PVT inputs on the LFP spectrogram. *Right*, Population data for theta (θ) band power changes provoked by inhibition of VLM_GABA_-to-PVT inputs (green) relative to pre-stim (black) in *Hight Theta* STIM trials (n = 7 animals; Wilcoxon matched-pairs test). *P < 0.05; **P < 0.01; ***P < 0.001. (Statistics in **Extended Data Table 1**).

To search for a neuronal pathway through which the VLM_GABA_ signal can reach PFC, we mapped the brain-wide efferentation of VLM_GABA_ neurons. First, we confirmed that cortical regions in general, including the PFC, receive no or negligible direct innervation from the VLM_GABA_ neurons (Extended Data Fig. 2). Conversely, widespread VLM_GABA_ inputs were observed subcortically, in the brainstem and hypothalamus, with dense axonal arborization also observed in the PVT. A quantitative analysis revealed that VLM_GABA_ neurons topographically projected to the PVT with the strongest input localized to the middle-posterior region (Fig. 1c,d), where PFC-projecting PVT neurons were identified by retrograde labeling (Fig. 1e,f). These suggest that these PVT neurons may represent a key relay linking medullary REM-sleep promoting activity to cortical networks.

To demonstrate functional involvement of the PVT in REM sleep, we next examined the activity of PVT neurons across sleep-wake cycles. To target PVT neurons, we used the calretinin (CR, ‘*Calb2*’)-Cre mice, as the majority of PFC-projecting PVT cells express calretinin^25^. We injected the PVT of CR-Cre mice with a Cre-dependent recombinant AAV expressing ChR2. Three weeks later, we implanted optrode-tetrode configured wire bundles surrounding an optical fiber to record the activity of optogenetically identified PVT cells during natural home-cage sleep. Simultaneously, EEG and EMG electrodes were placed above the frontal cortex and in the neck muscle to classify brain states (awake, non-REM, REM) based on theta (4-8 Hz for REM) and delta (1-4 Hz for NREM) oscillations, together with EMG signals (Extended Data Fig. 1a and 3a).

Analysis of identified PVT cells (N = 34 cells, n = 3 mice; Extended Data Fig. 3b) revealed diverse state-dependent activity modulations (Extended Data Fig. 3c-e). At population level, PVT neurons fired at significantly higher rates during REM sleep compared to NREM sleep (Extended Data Fig. 3c).

The most pronounced increase in the PVT firing occurred at the NREM-to-REM transitions (Extended Data Fig. 3e,f) highlighting their REM-dependence activity pattern. Moreover, a negative correlation was also observed between baseline firing rate and REM-induced firing rate changes, indicating that neurons with lower baseline activity exhibited greater REM-associated activation (Extended Data Fig. 3d).

Next, we examined whether VLM_GABA_ signaling contributes to shaping the sleep stage-dependent activity of PVT neurons and, in turn, cortical oscillations. To do so, we virally expressed NpHR3.0 in VLM_GABA_ cells of vGAT-Cre mice (n = 7) and performed simultaneous in vivo extracellular recordings from PVT and PFC while optogenetically inhibiting VLM_GABA_-to-PVT terminals under anesthesia (Fig. 1g). This configuration allowed us to perform cell-type-selective thalamic recordings over extended periods. It also enabled us to differentiate local field potential signal (LFP) from superficial (layers 1-3) and deep (layers 5-6) PFC regions, given that sleep oscillations exhibit laminar profiles, with higher REM associated cortical activity in superficial layers^18,35–37^. Consistent with the heterogeneous sleep state-dependent activity of PVT cells (Extended Data Fig. 3c,d,g,h), inhibition of VLM_GABA_ input produced diverse firing rate changes in individual PVT cells, particularly during REM-like high-theta periods (N=95 cells, n = 7 mice; Fig. 1h). Notably, VLM_GABA_ modulated cells (PVT_MOD_; N = 68) exhibited lower baseline firing rates than non-modulated ones (PVT_MOD_, N = 68: 0.74±0.85 Hz; PVT_NON_, N = 27: 1.85±1.97 Hz), and only PVT_MOD_ neurons showed increased activity during the NREM-to-REM transitions (Fig. 1i,j and Extended Data Fig. 2a,b). Moreover, REM-associated firing rate changes correlated with baseline activity only in PVT_MOD_ but not in PVT_NON_ neurons (Fig. 1k) and only firing of PVT_MOD_ neurons was phase-locked to the ascending theta cycle, suggesting their involvement in promoting theta oscillations (Fig. 1l).

We also found that optogenetic silencing of VLM_GABA_ input, both non-selectively and specifically during high-theta states, disrupted ongoing theta oscillations in superficial but not in deep PFC layers (Fig. 1m and Extended Data Fig. 4c-j). Notably, REM-associated silencing exerted a stronger theta-suppressive effect in 6 out of 7 mice (Extended Data Fig. 4k). Altogether, these findings indicate that a subpopulation of PVT neurons is under REM-related VLM_GABA_ control and influences theta oscillations in a VLM_GABA_-dependent manner.

### PVT_→NAc_ neurons selectively mediate the REM sleep-promoting and theta-modulating effect of VLM_GABA_ neurons

A subset of low firing PVT neurons mediates VLM-driven theta oscillatory changes. The PVT-to-nucleus accumbens (PVT_→NAc_) and PVT-to-amygdala (PVT**_→_**_AMY_) pathways represent the two largest PVT subpopulations, each with distinct connectivity and functions^25,27,38,39^. To determine which subgroup is under VLM_GABA_ control and exerts a laminar-selective influence on cortical oscillation and REM sleep, we employed multiple approaches. First, we retrogradely labeled PVT**_→_**_NAc_ or PVT**_→_**_AMY_ neurons using rgAAV-eYFP while VLM_GABA_ cells were transduced with AAV carrying ChR2 in vGAT-Cre mice. After 8–10 weeks, we conducted acute slice electrophysiology on coronal slices from the middle-posterior PVT (Fig. 2a,b). Optogenetic activation of VLM_GABA_-to-PVT fibers evoked GABA_A_-mediated inhibitory postsynaptic potentials (eIPSCs) in both PVT**_→_**_NAc_ and PVT**_→_**_AMY_ neurons (Fig. 2c). However, eIPSCs amplitudes were significantly larger than spontaneous IPSCs in PVT**_→_**_NAc_ but not PVT**_→_**_AMY_ neurons (Fig. 2d). Additionally, PVT**_→_**_NAc_ neurons exhibited stronger and more reliable inhibitory responses than PVT**_→_**_AMY_ neurons across all stimulation frequencies (Fig. 2e,f). While VLM_GABA_-to-PVT**_→_**_AMY_ synapses displayed short-term depression at 5/10/20 Hz, VLM_GABA_-to-PVT**_→_**_NAc_ synapses maintained stable transmission (Fig. 2e,f), suggesting that PVT**_→_**_NAc_ neurons more efficiently relay VLM_GABA_ inputs.

**Fig. 2:**
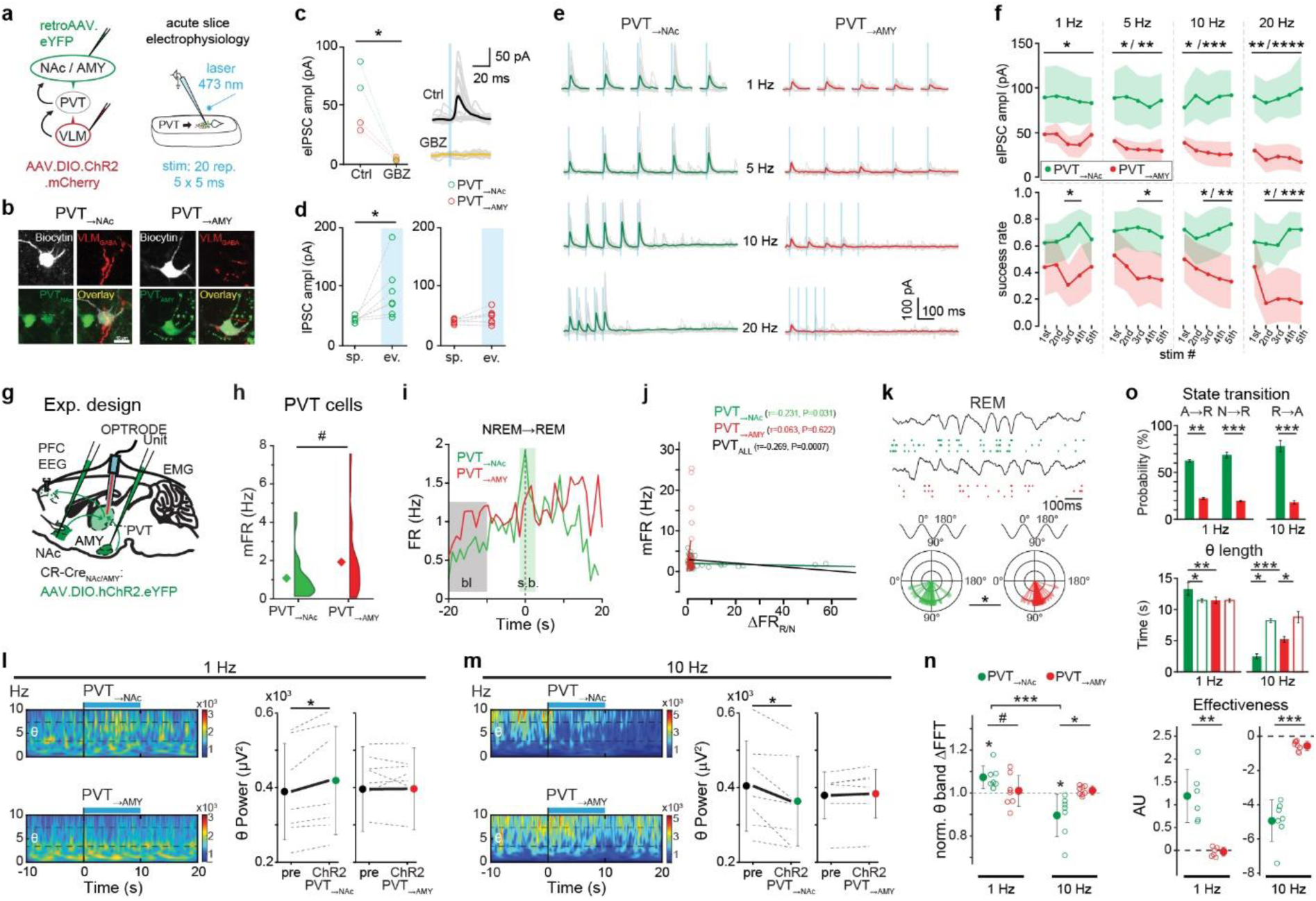
PVT_→NAc_ neurons mediate the impact of VLM_GABA_ on cortical theta oscillation and REM sleep. **a,** Experimental design for slice electrophysiology in PVT of vGAT-Cre mice examining VLM_GABA_ inputs onto projection-specific PVT cells. **b,** Representative high-magnification Z-stack confocal images (3 μm total in depth) showing biocytin-filled (white), PVT**_→_**_NAc_ (green; *left*) and PVT**_→_**_AMY_ (green; *right*) cells contacted by VLM_GABA_ inputs (red). **c,** *Left,*optogenetically evoked GABAergic IPSCs (eIPSC; Ctrl) in PVT cells (N = 4) were completely abolished by gabazine (GBZ; orange). *Right,* example traces of a PVT_→NAc_ cell in Ctrl (black) and GBZ (orange) conditions (Student’s t-test). **d**, Population data for spontaneous and evoked IPSCs of PVT**_→_**_NAc_ (green) and PVT**_→_**_AMY_ cells (red; N = 7-7; Student’s t-test). **e,** Example traces of a PVT**_→_**_NAc_ and PVT**_→_**_AMY_ cell for frequency-dependent eIPSCs. **f,** Population data for eIPSC amplitude and success rate (One-Way ANOVA, Bonferroni’s post hoc comparisons). **g,** Experimental design for in vivo recordings of PVT**_→_**_NAc_ or PVT**_→_**_AMY_ cells during natural sleep. **h,** Differences in mean firing rate (mFR) of PVT**_→_**_NAc_ (N = 25 cells; n = 7 animals) or PVT**_→_**_AMY_ cells (N = 36 cells; n = 8 animals; Mann-Whitney U test). **i,** Elevation in mean firing rate at NREM-to-REM transition for PVT**_→_**_NAc_ (*left*) but not PVT**_→_**_AMY_ cells (*right*; Wilcoxon matched-pairs test). Pink color, 1 s bins with significant elevation (s.b.) compared to baseline (bl, grey). **j,** Negative correlation between mean firing rate (mFR) and REM induced firing rate changes (R/N) restricted to PVT**_→_**_NAc_ cells Kendall’s TAU Correlations). **k,** Distinct phase-locked firing of PVT**_→_**_NAc_ (green) and PVT**_→_**_AMY_ cells (red) during REM-associated theta oscillation (Pearson’s χ2 test). *Top,* example unit activity of a PVT**_→_**_NAc_ (green) and PVT**_→_**_AMY_ cell (red). *Bottom,* vector diagram of population data. **l,m,** Frequency-dependent bidirectional effects of optogenetic stimulation of PVT**_→_**_NAc_ and PVT**_→_**_AMY_ cells on REM-associated theta oscillation indicated by wavelets (**l**, **m**, *left*) and quantification of theta band power changes (**l,m**, *right*). **n,** Summary data for the impact of frequency-dependent optogenetic stimulation on theta band power (n = 7-8 mice; Wilcoxon matched-pairs test). **o,** Population data for optogenetically evoked state transitions (*top*), duration of REM episodes (*middle*) and the calculated effectiveness to change REM sleep (*bottom*). ^#^P < 0.1; *P < 0.05; **P < 0.01; ***P < 0.001; ****P < 0.0001. (Statistics in **Extended Data Table 1**).

To confirm the selective involvement of PVT**_→_**_NAc_ neurons in REM sleep, we used retrograde labeling^25^ in CR-Cre mice to express ChR2 in either PVT**_→_**_NAc_ or PVT**_→_**_AMY_ neurons (Extended Data Fig. 5a-g), and recorded their firing rates across natural sleep stages (Fig. 2g and Extended Data Fig. 6a,b). PVT**_→_**_NAc_ neurons had lower baseline activity and stronger REM modulation compared to PVT**_→_**_AMY_ neurons (Fig. 2h and Extended Data Fig. 6b,c). Only PVT**_→_**_NAc_ neurons increased their firing at NREM-to-REM transitions (Fig. 2i and Extended Data Fig. 6d) and showed a negative correlation between baseline activity and REM-induced firing changes (Fig. 2j). Furthermore, PVT**_→_**_NAc_ firing was phase-locked to the ascending theta cycle, whereas PVT**_→_**_AMY_ firing aligned with the descending phase, suggesting a PVT**_→_**_NAc_-mediated role in theta oscillations (Fig. 2k) resembling VLM_GABA_-modulated PVT neurons (Fig. 1k,l). These findings establish PVT**_→_**_NAc_ as the key PVT

To directly asses the causal role of PVT**_→_**_NAc_ and PVT**_→_**_AMY_ neurons in REM sleep, we optogenetically activated these neurons at two different frequencies during natural sleep. Since PVT**_→_**_NAc_ activity naturally rises to ~1 Hz before NREM-to-REM transitions (Fig. 2i) and maintains this rate throughout REM bouts, we applied a 1 Hz stimulation protocol. In addition, a 10 Hz protocol was also carried out as activity of PVT cells were shown to reach this frequency^25^. Optogenetic activation of PVT**_→_**_NAc_ activation at 1 Hz (n = 7 mice) enhanced REM sleep, increasing theta band power and REM episode duration (Fig. 2l,n and Extended Data Fig. 6e,g). In contrast, 10 Hz PVT**_→_**_NAc_ activation suppressed REM-associated theta oscillations (Fig. 2m,n, and Extended Data Fig. 6f,h). Furthermore, PVT**_→_**_NAc_ activations triggered sleep-stage transitions with greater probability and efficacy and produced more pronounced REM disruptions, compared to PVT**_→_**_AMY_ (n = 8) or control PVT_YFP_ stimulations (n =3; Fig. 2l-o and Extended Data Fig. 6e-j). Together, these results indicate that PVT**_→_**_NAc_ neurons exert bidirectional control over REM sleep in an activity-dependent manner.

### PVT_→NAc_ bidirectionally modulate the VLM_GABA_ effects on theta oscillation

The previous results collectively suggest that PVT**_→_**_NAc_ is able to maintain or suppress the VLM_GABA_ input-mediated effects on theta oscillation. To directly verify this, PVT_CR+_ and VLM_GABA_ were transduced with AAVs carrying ChR2 in CR-Cre/vGAT-FLPO mice. After recovery, antidromic activation of PVT PVT_→NAc_ and orthodromic activation of VLM_GABA_ neurons were simultaneously applied while thalamic neuronal and cortical LFP activities were monitored (Fig. 3a). Optogenetic activation of VLM_GABA_ selectively altered the firing rate of the photo-identified PVT_→NAc_ but not of the un-identified PVT neurons (PVT_UN_; Fig. 3b). Notably, VLM_GABA_ activation increased PVT_→NAc_ firing to ~1 Hz. This elevation was further enchanced when VLM_GABA_ stimulation was paired with 1 Hz PVT_→NAc_ activation, and increased even more when it was combined with 10 Hz thalamic stimulation (Fig. 3c and Extended Data Fig. 7). In addition to changes in mean firing rate, PVT_→NAc_ neurons shifted from phasic to tonic firing modes when 1 Hz thalamic stimulation was replaced by 10 Hz stimulation together with VLM_GABA_ activation (Fig. 3d), as reflected by the interspike interval distribution (Fig. 3e).

**Fig. 3:**
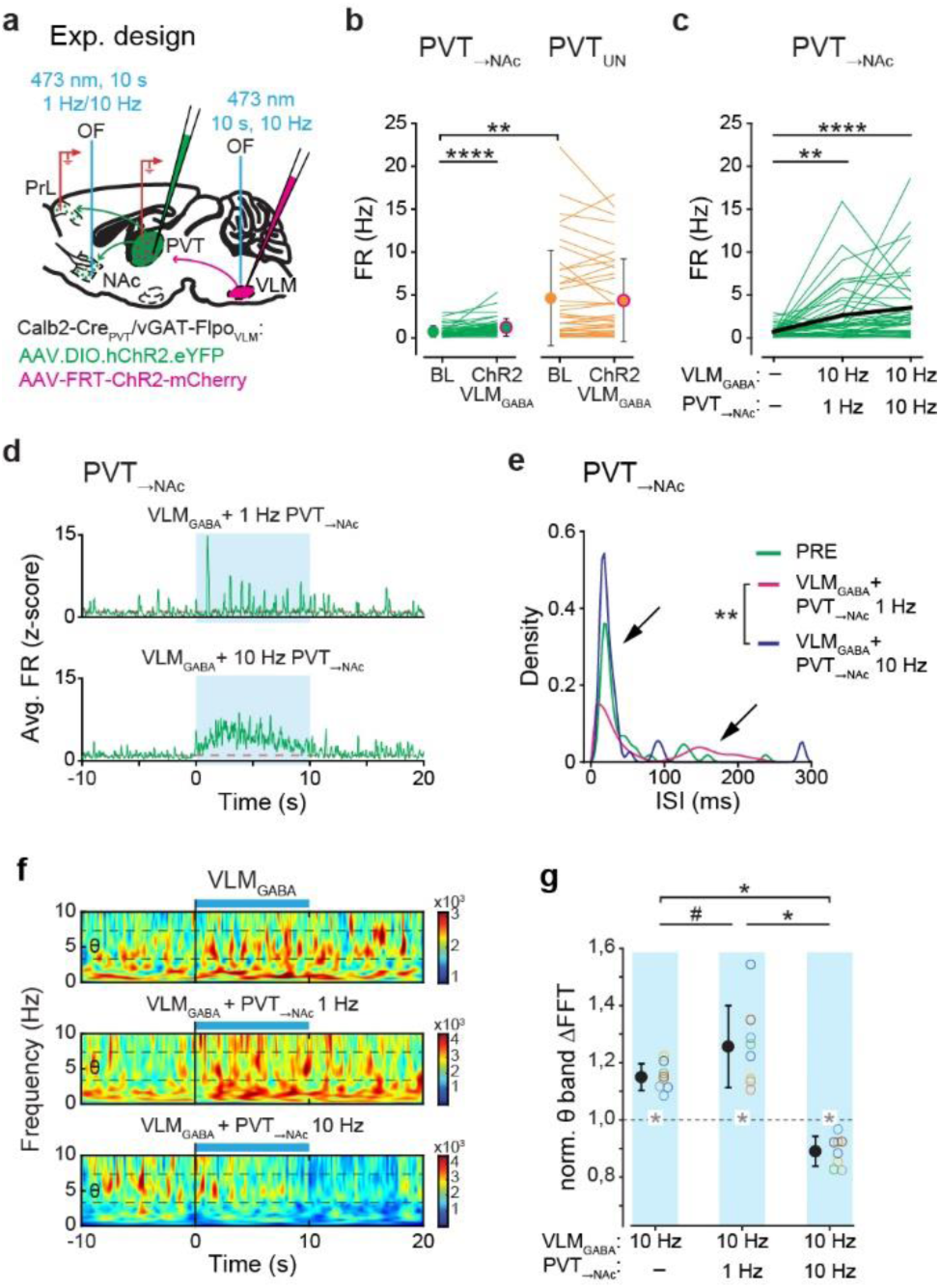
PVT_→NAc_ bidirectionally modulate the VLM_GABA_ effect on theta oscillation in activity- and firing mode-dependent manner. **a,** Experimental design for investigating the impact of VLM_GABA_ and PVT_→NAc_ optogenetic co-activation. **b,** Baseline (BL) and evoked PVT activity induced by stimulation of VLM_GABA_ neurons clustered by responsiveness to antidromic phototagging from NAc (PVT_→NAc_; N=41 cells) vs. untagged cells (PVT_UN_,N= 39 cells; n=8 mice; Wilcoxon matched-pairs test for BL vs. VLM_ChR2_; Mann-Whitney U test for PVT_→NAc_ vs. PVT_UN_). **c,** Baseline and evoked PVT_→NAc_ activity during paired stimulation of VLM_GABA_ and PVT_→NAc_ at 1 and 10 Hz (Wilcoxon matched-pairs test). **d,** Averaged activity changes of PVT_→NAc_ neurons evoked by VLM_GABA_ orthodromic and PVT_→NAc_ antidromic optogenetic co-activation indicated as z-scores. **e,** Inter-spike interval (ISI) distribution for PVT_→NAc_ neurons during pre-STIM and STIM periods (10 Hz VLM_GABA_ + 1 Hz PVT_→NAc_ and 10 Hz VLM_GABA_ + 10 Hz PVT_→NAc_). Note the multimodal ISI distribution (arrows) observed during 10 Hz VLM_GABA_ + 1 Hz PVT_→NAc_ stimulation (Kolmogorov-Smirnov test). **f,** Effect of the three stimulation protocols on the LFP spectrogram. **g,** Frequency-dependent, bidirectional effects of paired stimulation of VLM_GABA_ and PVT_→NAc_ on theta oscillation (Wilcoxon matched-pairs test). ^#^P < 0.1; *P < 0.05; **P < 0.01; ****P < 0.0001. (Statistics in **Extended Data Table 1**).

Meanwhile, this progressively elevated PVT_→NAc_ activity bidirectionally modulated the theta rhythm-promoting effects of VLM_GABA_ input: while 1 Hz PVT_→NAc_ activation maintained theta band power, 10 Hz thalamic stimulation suppressed it (Fig. 3f,g). These results confirm that, despite the ongoing VLM_GABA_ signaling, PVT_→NAc_ neurons can either enhance or suppress REM-associated PFC theta oscillations in an activity- and firing mode-dependent manner.

### Stress-induced REM sleep modulation by PVT_→NAc_ neurons

Having identified PVT_→NAc_ neurons as a key mediator of VLM_GABA_-to-PFC signaling in physiological regulation of REM sleep and the associated cortical theta oscillation, we next investigated whether this population also contributes to stress-induced modulation of REM sleep. Stress exerts bidirectional effects on REM sleep: while acute stress transiently enhances REM sleep as an adaptive response, chronic or intense stress suppresses it^40–43^. Thus, we employed a 4-day restraint stress paradigm, allowing us to monitor the activity of the same, optogenetically identified PVT cells during both acute and chronic stress as well as characteristics of REM sleep.

Following a habituation day (R0), mice underwent 3 days of restraint stress (R1-3), with day 1 (R1) representing acute phase and days 2-3 (R2-3) modelling chronic phase of the restraint stress (Fig. 4a). Throughout the experiment, frontal cortical EEG, EMG, and longitudinal thalamic single unit recordings were conducted during habituation/restraint and the subsequent 6-hour post restraint sleep periods (PRS0-3). Mice (n=6 mice) exhibited biphasic REM sleep modulation due to the restraint stress: REM sleep and the associated theta band power increased following acute phase, but were progressively suppressed upon repeated restraint, chronic stress. By the third day, both of them were significantly reduced compared to baseline (Fig. 4b and Extended Data Fig. 8a,b). Concurrently, PVT_→NAc_ neurons (N=11 cells; n=3 mice) showed increased firing during both the restraint and post-restraint sleep periods, with some cells reaching mean firing rates of 10 Hz, which response was absent in PVT_→AMY_ neurons (N=9 cells, n=3 mice; Fig. 4c,d and Extended Data Fig. 8c-e).

**Fig. 4:**
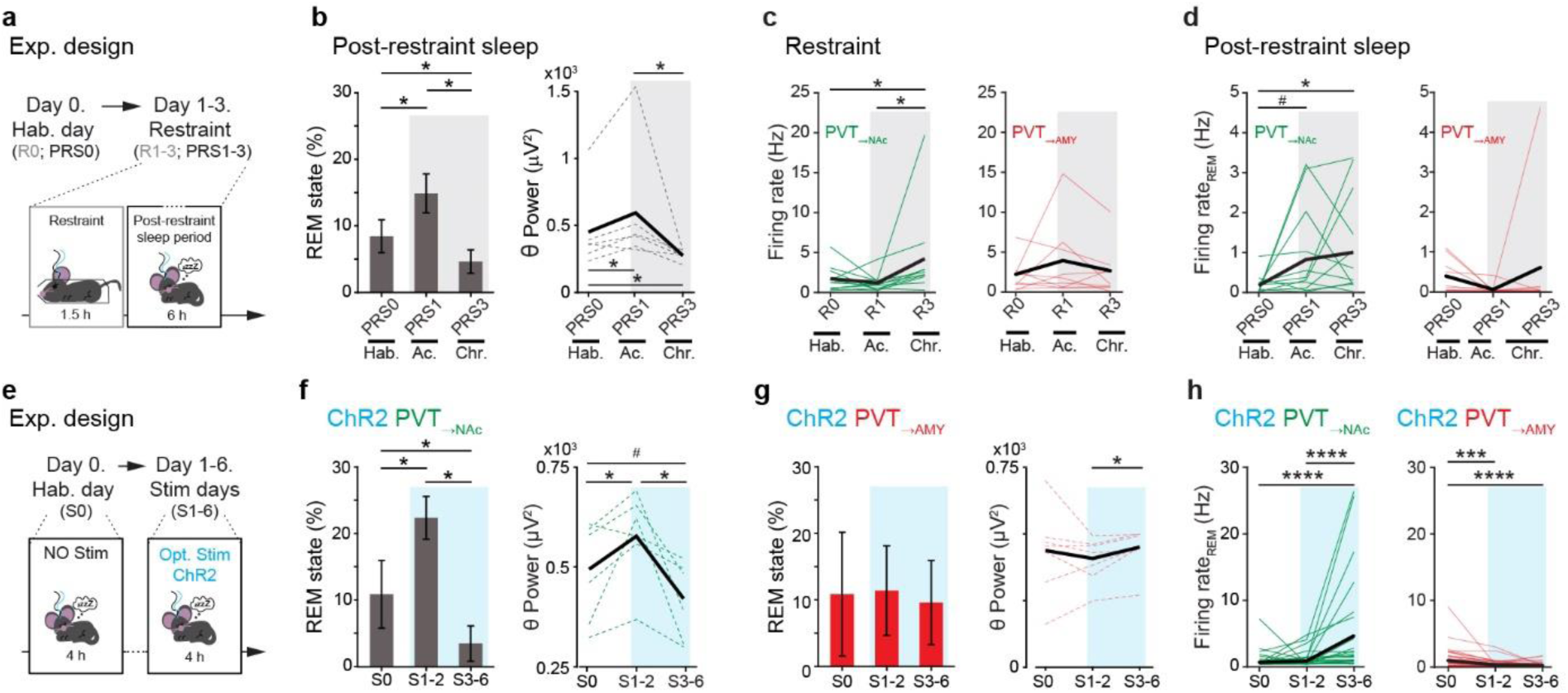
Activity changes of PVT_→NAc_ neurons are also responsible for stress-induced bidirectional modulation of REM sleep. **a,** Experimental design for investigating stress-dependent REM sleep alterations. **b,** Acute (Ac.) and chronic (Chr.) restraint induced changes in REM state during post-restraint sleep periods (PRS1-3) compared to the habituation day (Hab.; PRS0) indicated as proportion of REM sleep (*left*) and theta-band power (*right*) (n = 6 mice; Wilcoxon matched-pairs test). **c,d,** Population data for restraint-induced activity changes of PVT_→NAc_ (*left*; N = 11 cells) and PVT_→AMY_ cells (*right*; N = 9 cells) during restraint [R0-R3; (**c**)] and post-restraint REM sleep periods [PRS0-PRS3; (**d**)] (Repeated Measures ANOVA with post hoc LSD test). **e,** Experimental design for investigating the impacts of optogenetic activation of PVT_→NAc_ (n = 7 mice) and PVT_→AMY_ neurons (n = 8 mice) on REM sleep. **f,g,** Optogenetic stimulation (blue) of PVT_→NAc_ (**f**) but not PVT_→AMY_ (**g**) triggered REM state changes similar to those induced by acute and chronic restraint stress (Wilcoxon matched-pairs test). **h,** Population data indicating REM-sleep-related activity changes of PVT_→NAc_ (N = 25) and PVT_→AMY_ (N= 36) neurons during non-stim periods (Repeated Measures ANOVA with post hoc LSD test). *P < 0.05; **P < 0.01; ****P < 0.0001. (Statistics in **Extended Data Table 1**).

To establish a causal link between the progressive increase in PVT_→NAc_ activity and bidirectional REM sleep modulation, we optogenetically activated PVT_→NAc_ neurons (10 Hz, 10 s) during sleep across multiple days while monitoring frontal cortical EEG activity (n = 7 mice; Fig. 4e). Control groups included mice expressing ChR2 in PVT_→AMY_ neurons (n = 8) and YFP in PVT (n = 3).

Despite individual variability, PVT_→NAc_ stimulation produced bidirectional REM modulation, mirroring the effects of restraint stress: stimulation on the first two days (S1-2) increased REM sleep and theta band power, resembling the acute stress responses, whereas continued stimulation (S3-6) progressively suppressed both, similar to chronic stress (Fig. 6f and Extended Data Fig. 8h). PVT_→NAc_ firing also exhibited a progressive increase in non-stimulated periods, paralleling the impact of restraint stress (Fig. 4h and Extended Data Fig. 8k). In contrast, stimulation of PVT_→AMY_ and YFP expressing mice showed no REM sleep alterations (Fig. 4g and Extended Data Fig. 8i,l) meanwhile the activity of PVT_→AMY_ cells did not increase but decreased (Fig. 4h). The comparable REM sleep dynamics induced by restraint stress and PVT_→NAc_ activation (Fig. 6, b,f and Extended Data Fig. 8j) indicate that progressive PVT_→NAc_ recruitment mediates stress-dependent REM sleep plasticity.

We also aimed to determine whether the PVT_→NAc_ is embedded into a brain-wide network collecting the necessary stress and other theta-rhythm signals to shape sleep architecture in a stress-dependent manner. Rabies-mediated monosynaptic retrograde tracing revealed that PVT_→NAc_ neurons receive a broader array of inputs than PVT_→AMY_, including projections from stress-sensitive and REM-regulating hypothalamic, mesencephalic and brainstem nuclei (Extended Data Fig. 9a-h). These findings position PVT_→NAc_ as a central hub for integrating stress and REM rhythm signals and modulating theta oscillation during sleep, accordingly.

### Activity-dependent effects of PVT_→NAc_ on PFC circuit

Next, we examined the impacts of the presently described medullo-thalamic connection on cortical microcircuit dynamics responsible for REM-associated theta oscillation^18,44^. During acute *in vivo* extracellular LFP recordings, we also monitored single-unit activity of principal neurons (PN) as well as fast-spiking (FS) and non-fast-spiking (NFS) interneurons in PFC, classified based on their electrophysiological properties (Extended Data Fig. 10a-d). In parallel with PVT activity and cortical theta oscillation changes (Fig. 1), prefrontal cortical interneurons (IN) and PN were also modulated by the inhibition of VLM_GABA_-to-PVT terminals. A larger proportion of IN and PN exhibited activity changes when optogenetic silencing occurred during REM-like high theta states compared to all trials Our further analysis revealed that FS neurons decreased while PN and NFS neurons increased their activity upon inhibition of VLM_GABA_-to-PVT inputs, coinciding with the reduction of theta oscillation (Extended Data Fig. 10f-i). These findings suggest that the VLM_GABA_ inputs exert an influence on PVT, which in turn, shape the REM-associated theta rhythm via the PFC circuit.

To find out whether selective activation of PVT_→NAc_ – which bidirectionally modulates REM sleep and the associated theta oscillations (Fig. 2) – can drive similar changes in prefrontal microcircuit^18,44^, we further dissected the PVT-PFC network. First, we compared the thalamic axonal distribution of PVT_→NAc_ and PVT_→AMY_ cells in the PFC. Analyzing the retro-anterogradely labeled thalamic axonal patterns (see *Methods*), PVT_→NAc_ neurons were found to preferentially target the superficial layers, while PVT_→AMY_ innervated the deep layers (Fig. 5a,b and Extended Data Fig. 11). Next, we investigated the existence of laminar-specific impact of PVT_→NAc_ and PVT_→AMY_ activation on cellular cortical activity. Antidromic activation of PVT_→NAc_ and PVT_→AMY_ neurons was robust (Extended Data Fig. 12) and induced distinct, layer-specific effects in prefrontal cortical single-unit activity, consistent with their projection patterns. PVT_→NAc_ activation engaged a larger fraction of PFC neurons and resulted in a better signal-to noise ratio in superficial layers compared to PVT_→AMY_ (Fig. 5c-h).

**Fig. 5:**
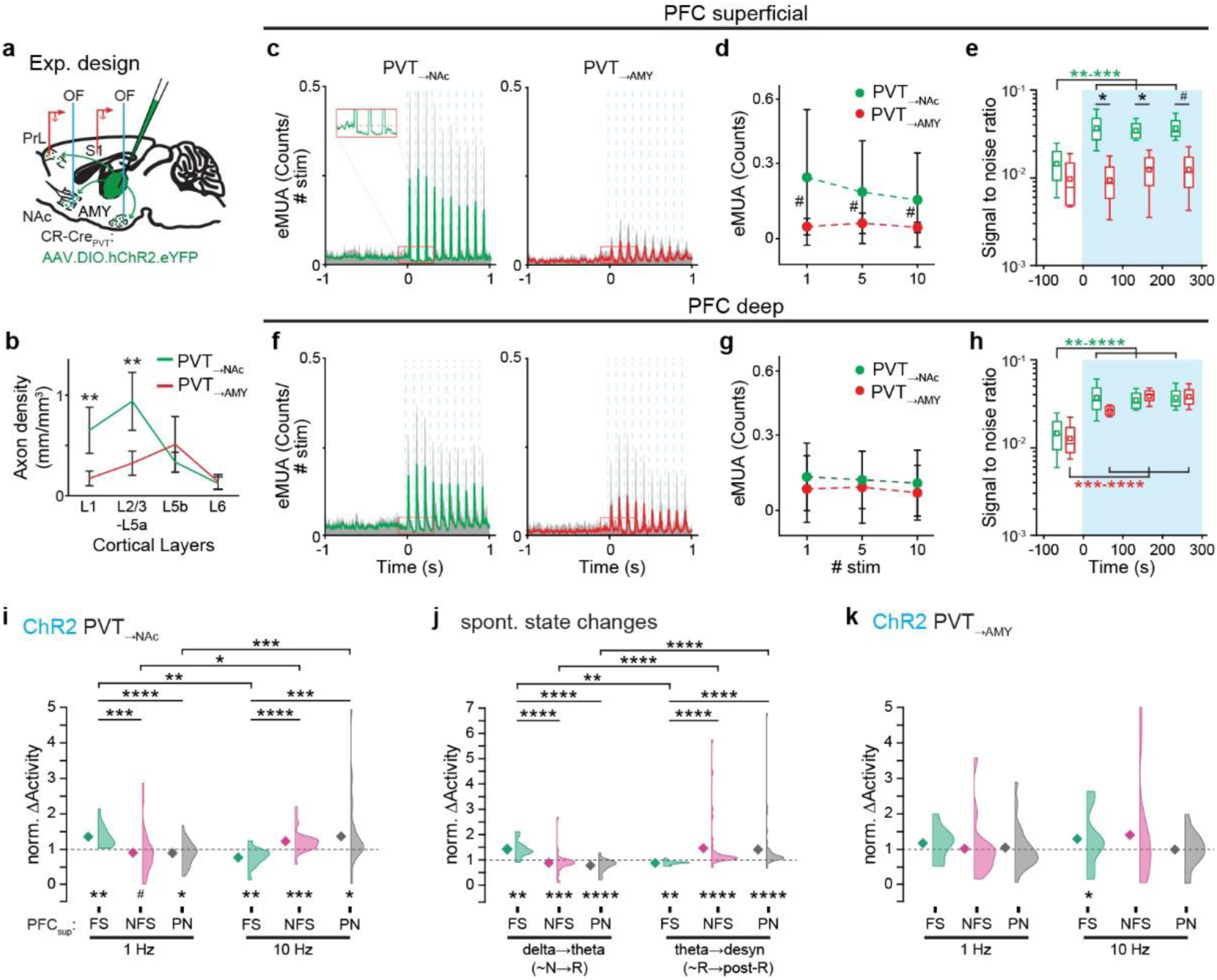
Frequency-dependent and layer-specific effects of PVT_→NAc_ on PFC circuits underlying modulation of cortical theta oscillation. **a,** Experimental design for investigating the effects of PVT_→NAc_ or PVT_→AMY_ activation on PFC circuits. **b,** Distinct, layer-specific distribution of PVT_→NAc_ and PVT_→AMY_ inputs in PFC (One-way ANOVA with post hoc LSD test). **c-h,** Distinct, layer-specific effects of PVT_→NAc_ and PVT_→AMY_ activation in PFC. Representative cases for evoked multiunit activity (eMUA) in the superficial (**c**) and deep layers of PFC (**f**); population data for eMUA (**a,g**; n= 4 mice; Wilcoxon matched-pairs test); population data for signal-to-noise ratio of eMUA (**e,h**; n=4 mice; Repeated Measures ANOVA with post hoc LSD test). **i,** Opposite effects of PFC FS (N=13 cells), NFS (N= 33 cells) and PN (N= 37 cells) triggered by 1 Hz and 10 Hz activation of PVT_→NAc_ (n=7 mice; Wilcoxon matched-pairs test for baseline vs 1 Hz vs. 10 Hz; Mann-Whitney U test for FS vs. NFS vs PN). **j,** Similar shifts in PFC FS, NFS and PN activation at spontaneous state changes: from high delta periods to high theta periods [similar to NREM (N) → REM (R)] and from high theta periods to desynchronized state [similar to REM (R) → post-REM (R)] (n=7 mice; Wilcoxon matched-pairs test for delta→theta vs. theta→desyn; Mann-Whitney U test for FS vs. NFS vs. PN). **k,** PVT_→AMY_ stimulation did not elicit state-dependent activation of PFC FS, NFS and PN (n=7 mice; Wilcoxon matched-pairs test for baseline vs 1 Hz vs 10 Hz; Kruskal-Wallis ANOVA for FS vs. NFS vs. PN). *P < 0.05; **P < 0.01; ***P < 0.001; ****P < 0.0001. (Statistics in **Extended Data Table 1**).

Notably, in anesthetized condition, 1 Hz and 10 Hz stimulation of PVT_→NAc_ (Extended Data Fig. 13a,b) elicited opposing changes in the superficial PFC theta power (increased and decreased; respectively) mirroring the effects observed in natural sleep conditions (Fig. 2). Similar LFP dynamics were detected in the primary somatosensory cortex (Extended Data Fig. 13c,d), indicating that PVT_→NAc_ — but not PVT_→AMY_ — can orchestrate global changes in cortical theta oscillation.

When 1 Hz PVT_→NAc_ activation enhanced theta power in PFC, FS neuron activity increased, while NFS and PN activity decreased. The opposite occurred with 10 Hz stimulation, which suppressed theta power (Fig. 5i and Extended Data Fig. 13e). Similar shifts in PFC microcircuit were observed during spontaneous sleep state transitions: from low-to-high theta (as in NREM-to-REM) and high-theta-to-desynchronized states (as in REM-to wake transitions; Fig. 5j and Extended Data Fig. 13f). Altogether, these findings indicate that PVT_→NAc_, but not PVT_→AMY_ (Fig. 5k and Extended Data Fig. 13g) selectively shapes the superficial PFC microcircuit dynamics underlying theta oscillation in an activity-dependent and stress-dependent manner (Fig. 6).

**Fig. 6:**
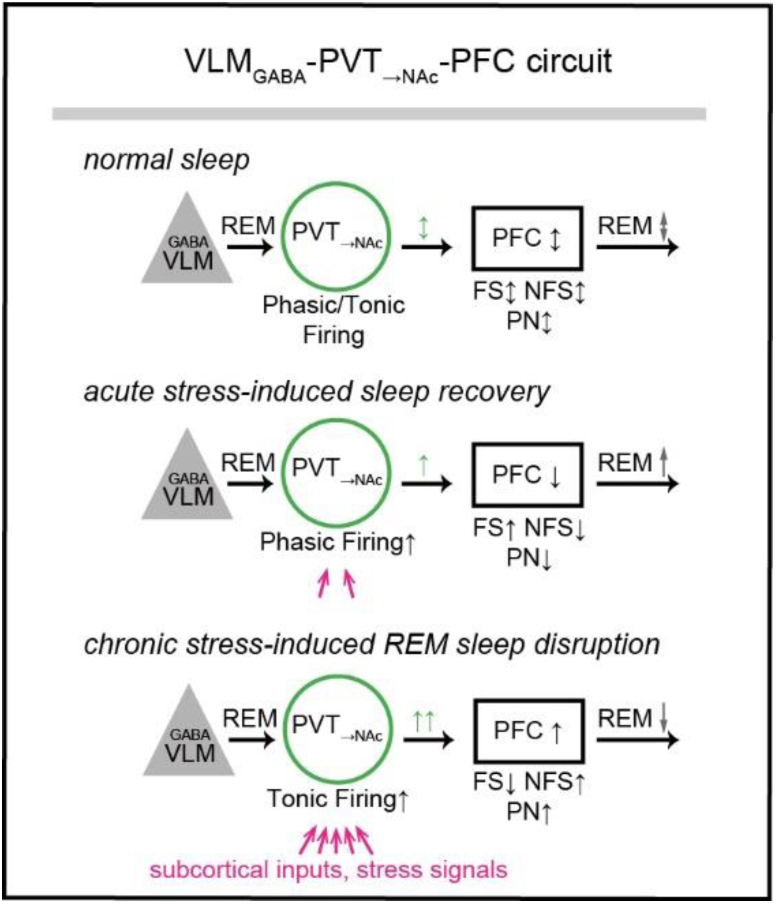
Circuit diagram for PVT_→NAc_ activity-mediated REM sleep alterations in normal and stressful conditions.

## Discussion

Establishing a neuronal network linking REM generator brainstem to PFC has broad implications for understanding how REM sleep architecture adapts to homeostatic needs or stress experiences. Our findings reveal a convergence of inputs that regulate REM sleep rhythm, such as from the REM rhythm-generating VLM_GABA_ neurons^14,15^ (Figs. 1-3), and from stress-signal carrying brain-wide pathways^45,46^ onto the PVT_→NAc_ population (Extended Data Fig. 9). In an activity- and firing mode-dependent manner, PVT_→NAc_ maintains or suppresses the effect of VLM_GABA_ signaling and drives shifts in the excitatory/inhibitory (E/I) balance within the PFC, which, in turn, are manifested as bidirectional changes in REM-associated theta oscillation (Figs. 3,5 and 6). We further demonstrate that PVT_→NAc_ activity increases progressively with prolonged stress exposure, mediating dual roles of stress in regulating REM sleep: acute stress results in a REM rebound, whereas repeated stress elevates PVT_→NAc_ activity and and disrupts REM sleep (Fig. 4).

VLM_GABA_ inputs to the PVT_→NAc_ were shown to play an important role in fine-tuning of REM sleep-associated theta oscillation. Short (5 ms) activation of VLM_GABA_ neurons exerts an inhibitory effect on individual PVT_→NAc_ cell by eliciting IPSCs (Fig. 2). However, longer (10 s) medullary inhibition caused facilitation at population level (Fig. 3). Prolonged inhibition can gradually raise intracellular chloride levels shifting the chloride equilibrium potential toward more depolarized values and thereby biasing the membrane potential closer to the thalamic firing threshold. Under such conditions, the same inhibitory input can become functionally excitatory in hippocampal and hypothalamic cells^47–49^, a scenario that may also be present in PVT_→NAc_ cells explaining their increased firing upon longer medullary inhibition (Fig. 2). This moderate enhancement in PVT_→NAc_ activity biases the system toward REM sleep transition.

In addition to the VLM_GABA_ input, the PVT_→NAc_ integrates a range of subcortical inputs essential for fine-tuning REM rhythm [such as the locus coeruleus^34,50^, lateral hypothalamus^51^, tegmental/pontine area^52^ and preoptic area^53^], regulating homeostatic needs^45^ as well as providing adequate behavioral responses to stress^30–33^ (Extended Data Fig. 9). The convergence of all of these inputs can further modulate PVT_→NAc_ firing mode and activity, which, in turn, re-evaluates REM rhythm transition to the PFC and modulates REM sleep theta oscillation, accordingly (Fig. 3).

The emergence of REM sleep-coupled theta oscillation in the cortex highly depends on the cortical E/I balance^18,37,44^. Our data highlight that PVT_→NAc_-PFC input acts as a critical modulator capable of altering PFC inhibitory circuits to adaptively promote or suppress cortical theta oscillation, in an activity-dependent manner (Fig 5). During acute (mild) stress or when homeostatic REM sleep pressure increases, PVT_→NAc_ neurons likely receive both moderately enhanced stress- or homeostasis-related excitatory input and prolonged VLM-mediated inhibition. This combination increases PVT→NAc activity engaging fast-spiking (FS) interneuron–mediated feedforward inhibition and shifting the cortical E/I balance toward REM sleep promotion^18,54,55^. Conversely, stronger excitatory inputs due to chronic stress can cause over-activation and tonic firing of PVT_→NAc_ neurons, shifting the E/I balance toward non-FS-based inhibition and REM sleep suppression. Consistently, we showed that acute 1 Hz and 10 Hz optogenetic stimulation of PVT_→NAc_ recapitulated homoeostatic changes in theta oscillation, whereas chronic stimulations mimicked stress-induced alterations of REM sleep. Repeated 10 Hz activation of PVT_→NAc_ neurons for 1-2 days resulted in facilitation of REM sleep theta oscillation similar to acute stress, while prolonged stimulation (up to 6 days) strongly reduced it similar to chronic stress. These findings provide causal evidence that establishes PVT_→NAc_ activity as an important mechanism in fine-tuning of REM sleep and its associated theta oscillation in accordance with physiological and behavioral states.

Notably, stressed-triggered thalamic hyperactivity is persistent, causing REM sleep facilitation or reduction for hours or even days, in contrast to the transient thalamic activity increase observed at natural NREM-to-REM transitions. Beyond fast glutamatergic transmissions, these long-lasting changes in sleep likely depend on slower neuromodulatory signaling mechanisms. One candidate is nitric oxide (NO), whose levels has been shown to be suppressed by acute stress while elevated by repeated stress^56^. As cortical NO level can influence E/I balance^57^ and that the PVT_→NAc_ cells express NO synthetizing enzymes (Extended Data Fig. 14), thalamic NO release may contribute to activity-dependent shifts in PFC inhibitory circuits underlying REM sleep modulation. Similarly, acute stress has been shown to elevate corticotrophin-releasing hormone (CRH) contributing to REM sleep rebound^58–61^. Given that PVT_→NAc_ cells also express CRH^62^, their stress-induced activation and CRH release may also influence cortical activity over longer timescales. Together, NO and CRH signaling in PVT→NAc neurons may mediate the long-term impact of stress on REM sleep regulation.

Encoding stress signals and shaping adaptive stress responses are widely distributed across forebrain circuits, including the NAc and PFC^63–67^. Recently, PVT neurons, particularly those with accumbal or cortical collaterals, have been shown to modulate stress-induced and motivational behaviors^27,31,68–70^. By identifying the sleep rhythm regulatory role of PVT_→NAc_ cells, which co-project to both forebrain regions, this thalamic cell group may serve as a shared mechanism linking REM sleep expression with stress-coping forebrain networks, thereby coordinating REM sleep regulation with its associated functions.

The laminar-specific cortical connectivity of PVT_→NAc_ neurons (Fig. 5) is also highly dedicated for REM sleep theta modulation, as these preferentially target the superficial layers. These cortical layers possess a unique role in REM sleep, supported by strong feed-forward and feed-back inhibitory mechanisms^37,71–74^. Cortical FS neurons form dense perisomatic-targeting and axo-axonic connections with neighbouring pyramidal cells, mediating fast local feed-forward inhibition, that is particularly critical for REM initiation and maintenance within superficial layers^18,36,37,44,75,76^. In contrast, NFS neurons provide a feed-back inhibition, reducing REM-associated theta activity and promoting slow wave sleep^77^. In addition, layer 2/3, but not layer 5, pyramidal neurons in the retrosplenial cortex are strongly activated during REM stage, indicating that superficial principal cells are indispensable for REM sleep regulations^37^. As shifts in cortical inhibition underlying REM sleep kinetics are primarily affected by PVT_→NAc_ neurons, distinguishing them from neighbouring PVT_→AMY_ cells, PVT_→NAc_ neurons are ideal to fulfil the role of ‘conductor’ in state-dependent regulation of REM-related cortical dynamics.

Taken together, their activity dynamics and input-output organization uniquely position PVT_→NAc_ neurons to modulate cortical REM sleep in a homeostatic and stress-dependent manner, exerting opposite effects on theta oscillations via superficial cortical inhibitory circuits. As homeostatic E/I balance-mediated REM sleep is essential for PFC-dependent functions^78–80^, stress-driven neuronal reorganization – modulated by PVT_→NAc_ activity – may lead to persistent changes within frontal cortical structure and in REM-associated cognitive performance^3,72,81^, and ultimately, contribute to the development of psychiatric disorders^82–85^.

## Methods

### Animals

Adult (2–5 months old female and male) CR (*Calb2*)-Cre (RRID: IMSR_JAX:010774), vGAT (*Slc32a1*)-Cre and -FLPO (from The Jackson Laboratory) mice were used. We also crossbred mice to have CR-Cre/vGAT-Flpo mice for the experiments. Female mice were used only for the anatomical investigations. Animals from the same gender were group-housed in a humidity- and temperature-controlled environment. Animals were kept under standard laboratory conditions on a 12 h light/12 h dark cycle with food and water available ad libitum. The Regional and Institutional Committee of the Research Centre for Natural Sciences, the Institute of Experimental Medicine and the National Animal Research Authorities of Hungary (PE/EA/00080-4/2023) approved all procedures. After all surgeries, animals received subcutaneous Rimadyl (Carprofen, 1.4 mg/kg).

### Stereotactic surgeries

#### Viral injection

Mice were anesthetized with ketamine–xylazine (5:1, 3× dilution, ketamine: 100 mg/kg; xylazine: 4 mg/kg) and injected with one of the following AAV vectors with Nanoliter Injector (WPI) at a rate of 1 nl s^−1^ (50-200 nL): AAV-Ef1a-DIO-ChR2-eYFP (Penn Vector Core; titer: 5 × 10^12^ to 1 × 10^13^ genome copies (GC)/ml), AAV5-EF1a-DIO-eYFP-WPRE-hGH (Penn Vector Core; titer: 5 × 1012 GC/ml), AAV-Ef1a-DIO-ChR2-mCherry (Penn Vector Core; titer: 5 × 10^12^ to 1 × 10^13^ GC/ml), AAV-DFO-ChR2-eYFP virus (titer: 1 × 10^13^ GC/ml), AAV5-EF1a-DIO-mCherry (UNC Vector Core; titer: 5.3 × 10^12^ GC/ml), AAV-Ef1a-DIO-eNpHR3.0-eYFP (Penn Vector Core; titer: 3 × 10^1^^3^ GC/ml), AAV-CAG-FLEXFRT-ChR2-mCherry (Addgene; titer: ≥ 7 × 10^12^ GC/ml) and AAVrg-CAG-GFP (Addgene; titer: 7 × 10^12^ GC/ml). The target regions are (with anterior–posterior, lateral and dorsal–ventral coordinates): dorsal midline thalamus (paraventricular thalamus, PVT; −1-1.2/±0/3.2–2.8 mm (n=35 animals); amygdala (AMY; −1.8/±3.4/-3.7 mm) (n=18 animals); nucleus accumbens (NAc; +1.4/±1.1/–3.9–4.2 mm) (n=16 animals) and ventrolateral medulla (VLM; −7.3/±1.4/– 5.3 mm) (n= 27 animals).

#### Rabies tracing

The tracing method has been used and validated earlier^86^. CR-Cre mice were injected with 100 nL of AAV2/8-hSynFLEX-TVA-p2A-eGFP-p2A-oG (4.5×10^12^ GC/mL) into the PVT. After 4 weeks, mice were injected with rabies(ΔG)-EnvA-mCherry (100 nL; 3.5×10^7^ GC/mL) into the NAc or AMY (N = 3-3). Mice were perfused after 9-10 days of survival. Both the rabies (mCherry) and the AAV (eGFP) signal were amplified by immunofluorescent staining procedure (see below).

### Immunohistochemistry

All anatomical data were obtained from immunohistochemically stained mouse brain slices. Firstly, the tissue blocks were cut with a VT1200S Vibratome (Leica) into 50 μm coronal sections. Free-floating brain sections were intensively washed (5 times for 10 min) with 0.1 M PB. During immunohistochemical processing, all antibodies were diluted in 0.1 M PB. For high-quality fluorescent labeling, sections were treated with a blocking solution containing 10% normal donkey serum (NDS, Sigma-Aldrich, S30-M) or normal goat serum (NGS, Vector, S-1000, RRID: AB_2336615) and 0.5% Triton-X (Sigma-Aldrich) for 30 min at room temperature (RT). In the case of electrode track analysis, sections were cryoprotected in 30% sucrose solution overnight, and freeze thawed over liquid nitrogen and then, treated with 10% normal donkey serum solution to keep the visibility of the DiI (1,1′-Dioctadecyl-3,3,3′,3′-tetramethylindocarbocyanine perchlorate; Santa Cruz Biotechnology).

Then, the sections were incubated in primary antibody solution at RT overnight or for 2–3 days (at 4 °C). The following primary antibodies were used: green fluorescent protein (GFP, chicken, Life Technology: A10262, RRID: AB_2534023; 1:2000), mCherry (rabbit, BioVision: 5993-100, RRID: AB_1975001; 1:2000), FoxP2 (mouse, Merck Millipore: MABE415, RRID: AB_2721039; 1:500; Invitrogen: MA5-31419, RRID: AB_2787055; 1:2000; rabbit, Abcam: ab16046, RRID: AB_2107107; 1:500), Calb1(rabbit, SWANT: CB38, RRID: AB_10000340; 1:2000; mouse, SWANT: 300, RRID: AB_10000347; 1:2000), PV (mouse, SWANT: PV235, RRID: AB_10000343; 1:2000), CR (mouse, SWANT: 6B3, RRID: AB_10000320; 1:2000), Ctip2 (rat, Abcam: ab18465, RRID: AB_2064130; 1:500), WFS1 (rabbit, Proteintech: 26995-1-AP; RRID: AB_2880717; 1:500) and NOS (mouse, Sigma: N2280; RRID: AB_260754; 1:300). After the primary antibody incubation, the sections were treated with the following secondary IgG antibodies for fluorescent staining (1:500; 2 h at RT): Alexa 488-conjugated goat anti-chicken (GACh-A488; Molecular Probes: A11039, RRID: AB_142924), Alexa 488-conjugated donkey anti-mouse (DAM-A488; Jackson: 715-545-150, RRID: AB_2340846), Alexa 555-conjugated donkey anti-rabbit (DAR-A555; Invitrogen: A31572, RRID: AB_162543), Cy3-conjugated donkey anti-mouse (DAM-CY3; Jackson: 715-165-151, RRID: AB_2340813), Cy3-conjugated donkey anti-rabbit (DAR-CY3; Jackson: 711-165-152, RRID:AB_2307443), Alexa 594-conjugated donkey anti-rabbit (DAR-A594; Molecular Probes: A21207, RRID: AB_141637), Alexa 647-conjugated donkey anti-rabbit (DAR-A647; Jackson: 711-605-152, RRID: AB_2492288). In some cases, staining was enhanced after primary antibody incubation with biotinylated secondary antibodies (1:300; 2 h at RT), which are the followings: biotinylated goat anti-rabbit IgG (bHAR; Vector Laboratories: BA-1100, RRID: AB_2336201), biotinylated goat anti-mouse IgG (bHAM; Vector Laboratories: BA-2000, RRID: AB_2313581), Elite Avidin-Biotin Complex (eABC; Vector Laboratories: PK-6100, RRID: AB_2336819), streptavidin-conjugated green fluorescent antibodies (1:1000-2000; 2 hr. at RT) (SA-A488; Jackson: 016-540-084, RRID: AB_2337249), streptavidin-conjugated red fluorescent antibodies (SA-CY3; Jackson: 016-160-084, RRID: AB_2337244), streptavidin-conjugated far red fluorescent antibodies (SA-A647; Jackson: 016-600-084, RRID: AB_2341101). All the fluorescent slices were mounted in Vectashield Mounting Medium (Vector Laboratories: H-1000, RRID: AB_2336789) or in Vectashield Mounting Medium with DAPI (Vector Laboratories: H-1200, RRID: AB_2336790).

Alternatively, we used DAB-Ni as a chromogen for Rabies. Sections were cryoprotected in 30% sucrose solution overnight, and freeze thawed over liquid nitrogen and then were treated with 1% H_2_O_2_ solution (10 min) for endogenous peroxide blocking. Next, the slices were incubated in 10% normal donkey serum (30 min, RT). After primary antibody incubation, slices were treated with biotinylated secondary antibodies (goat anti-rabbit or horse anti-goat IgG) and eABC (see above). To visualize the labeled cells, sections were developed with DAB-Ni. Sections were then dehydrated in xylol and mounted in DePex (Serva).

#### Identification of cortical layers

To differentiate the cortical layers in PrL and the primary somatosensory cortex (S1), we used combination of neurochemical markers^87^. The cortical layer 2/3 (L2/3) was identified by using Calb1 staining, while layer 6 (L6) was visualized with fork head box protein P2 (FoxP2) staining. Furthermore, for dissemination of the layer 5 were used Chicken ovalbumin upstream promoter transcription factor-interacting proteins 2 (Ctip2) and anti-Wolfram syndrome 1 (WFS1). The dorsoventral borders of the PrL cortex is determined by the parvalbumin (PV) immunostaining^88^.

#### Axon density analysis

We performed retro-anterograde viral labeling approach in CR-Cre mice to visualize PFC projections of PVT_→NAc_ and PVT_→AMY_ neurons as described earlier^25^. Shortly, AAV injection into NAc or AMY retrogadely-labeled CR+ neurons in PVT. If these PVT cells had cortical collaterals, the virus propagated in an anterograde fashion and visualized axons in PFC as well. These retro-anterograde viral labelings were also amplified and the slices were counterstained for the above mentioned markers to identify the PrL cortex layers and borders. Confocal Z-stack (slice number: 50; total thickness: 23.34-40.3 μm, PVT_→NAc_: 36.31±2.66 μm; PVT_→AMY_: 31.83±4.32 μm) images were taken with confocal microscope (Zeiss LSM 710; Zeiss ZEN 2010B SP1 Release version 6.0; Carl Zeiss Microimaging GmbH) a ×63 objective (10+x Plan Apochromat 10x/0.45 M27; 20x Plan Apochromat 20x/0.8 M27; 63x Plan Apochromat 63x/1.4 Oil DIC M27) at four different AP levels between Bregma +2.22 and +1.54 mm. The length of the retro-anterograde labelled CR+ axons were counted with a use of a custom-written Fiji macro^25^ in a given region of interest (available at https://github.com/baabek/Axon-density-analyzer-ImageJ-script.git). The macro calculated axon length for each stack and the total axon length was summed up for each layers and slices in each animal. Then, the relative axon density was counted for each layers and slices (total axon length / slice volume (ROI area * interval)), and finally the relative axon density was calculated for the animal (RAD = total axon length/total volume). Furthermore, we determined the total axon distribution by layers for each animal (layer RAD/all RAD).

To analyze the distribution of VLM_GABA_ axons in PVT, we used the same methods. The area of the PVT was determined by calretinin (CR) immunostaining^88^. The distribution of the anterogradely labelled VLM_GABA_ axons were counted between Bregma −0.3 and −2.1 mm in PVT.

### Chronic optrode implantations

Four weeks after AAV-Ef1a-DIO-ChR2-eYFP and AAV5-EF1a-DIO-eYFP virus injections into AMY or NAC in CR-Cre mice, four custom-fabricated tungsten tetrodes (12.5 μm in diameter, California Fine Wire) were implanted into the PVT (AP/ML/DV (−1/±0.6/–2.8 mm at 10°). The tungsten electrodes were attached with a multimode optic fiber (105-μm core diameter, NA = 0.22; Thorlabs) (200–300 μm left between the optic fiber and tetrode tips), with the help of a polyimide tube (0.008 ID, Neuralynx). The electrode wires were joined to an electrode interface board (EIB-16, Neuralynx) by gold electrode contact pins (Neuralynx). The ground, reference, EMG, EEG wires (Phoenix Wire Inc.) were soldered to the appropriate electrode interface board point. The tetrode impedances measured before the implantation at 1 kHz were kept between 240 and 750 kΩ (Intan Technologies). Ground and reference screws were implanted in parietal bone, respectively; an EMG wire was inserted into the neck muscle; an EEG screw was placed in frontal bone over PFC cortex. Finally, all pieces were secured onto the skull by multiple layers of dental acrylic (Paladur, Heraeus Kulzer) and shielded by a copper web. Mice were left for at least 4 days to recover and then, handled for several days.

### In vivo electrophysiology in anesthetized preparations

In vivo recordings were performed 4–12 weeks after the viral injections (AAV-Ef1a-DIO-ChR2-eYFP or AAV-CAG-FLEXFRT-ChR2-mCherry into the PVT or AAV-Ef1a-DIO-eNpHR3.0-eYFP into the VLM) from the prelimbic cortex, primer somatosensory cortex and the paraventricular thalamus in urethane anesthetized (20 m/m % dilution; 0.005ml/1g) CR-Cre or vGAT-Cre transgenic mice. The head of the animal was fixed in a stereotaxic frame; a screw was driven into the occipital or into the left parietal bone, which served as a reference electrode. LFP activity as well as the extracellular unit activities from PrL, PVT and S1 were monitored with silicon probes (A1x16-5mm-50-703, L16; A1x32-10mm-50-177, L32; A2x16-10mm-100-500-177, BL32; A1x32-5mm-50-177, L32; Neuronexus), stained with fluorescent DiI. The wideband neural data (0.1–7500 Hz) were amplified (gain: 192×) and digitalized at 20 kHz (Intan Technologies). The electrodes were lowered into PrL (AP/ML/DV +2/–2.5/–3.5 mm/60°), into S1 cortex (AP/ML/DV –1.2/+3.2/–1, –0.8 mm/20°) and into PVT (AP/ML/DV –1.1/+1/–3.2 mm/20°).

For the ‘antidromic–orthodromic’ activation^25^, the optic fibers were placed into the NAc and AMY, where the PVT_CR+_ fibers were activated, and the evoked LFP and unit responses were detected in PrL, S1 and PVT. Under these conditions, action potentials first traveled antidromically and, at a putative branching, could turn to orthodromic direction as well. The NAc and AMY did not contain CR+ cells, which projected back to the PVT, so the thalamic stimulation was due to direct antidromic activation. In the cases of PVT somatic and VLM_GABA_→PVT terminal manipulations, the optic fiber was placed above the PVT. For the VLM_GABA_ somatic stimulation, the optic fiber was placed above the VLM (AP/ML/DV – 7.2/+3.3/–5.6/20°). Optogenetic activation consisted of 473-nm blue-light pulses (1s or 10s, 1 or 10 Hz, ~10-15 mW, Laserglow Technologies). In the case of the 1 s long activations, the stimulus was repeated 30 times and the interstimulus intervals were 60 s, while the 10 s long stimulations were repeated 10 times with 250 or 300 s interstimulus intervals. The optogenetic inhibition with NphR3.0 consisted of 532-nm green-light pulses (10s, ~15 mW). The lasers were triggered by analog signals driven by a National Instruments acquisition board (USB-6353) and Matlab. Analog trigger pulses were also delivered to the recording system and registered parallel with neural data with zero latency. After recordings, animals were transcardially perfused, and coronal sections were cut from the paraformaldehyde-fixed brains. The position of the silicon probes were verified based on DiI labeling of the electrode track. For determination of the layer specific mapping, we used different neurochemical markers (see above). Mice with incorrect virus injections or electrode positions were excluded from the analysis (N=4 mice).

### In vivo electrophysiology of freely behaving mice

In vivo recordings from freely behaving CR-Cre mice were performed 4-5 days after the optrode implantation (into the PVT). EEG, EMG and extracellular PVT unit activities were monitored with optrodes and wire implantation (see previously). Of note, chronic EEG recording with frontal cortical screw electrodes and acute LFP recordings with multisite silicon probes resulted in comparable sleep oscillation fluctuations, similar duration of single theta band (~REM) episodes as well as similar theta band power values (Fig. S1). These findings indicate that both approaches are equally suitable for investigating the neuronal processes underlying REM-associated theta oscillations.

The wideband neural data (0.1–7500 Hz) were amplified (gain: 192×) and digitized at 20 kHz. Optogenetic tagging and activation consisted of 473-nm blue-light pulses (5 ms 0.5–1 Hz, for tagging; 10s 1 Hz; 10s 10 Hz; ~15 mW). The evoked unit responses were detected in PVT_CR+_, PVT_→NAc_ or PVT_→AMY_ neurons simultaneously with the EEG and EMG signal. The PVT_→NAc_ or PVT_→AMY_ neurons were transduced with ChR2 carrying AAV vectors using retro-anterograde viral injections. For optical identification, 5 ms long pulses were repeated 30 times, while the 10 s long 1 Hz and 10 Hz stimulations were repeated several times with random interstimulus intervals, which was minimum 300 s long. The behavior of mice was captured on video at 30 frames/s.

We measured the spontaneous sleep-wake transitions and sleep architecture of the PVT_→NAc_ (N= 6 mice) and the PVT_→AMY_ mice (N= 7 mice) for several days. Each day 10s 1 Hz and 10s 10 Hz stimulation were randomly applied.

#### Restraint stress

The spontaneous sleep-wake transitions and sleep architecture were measured in both light and dark phases (EEG, supplemented with EMG signal) and the activity of the PVT_→NAc_ and PVT_→AMY_ (N= 6 mice) cells. During the four-day experiment 19 units of PVT_→NAc_ and 20 units of PVT_→AMY_ neurons were followed. Only cells that could be followed through the entire stress paradigm were included in the further analysis (PVT_→NAc_, N = 11 and PVT_→AMY_, N = 9). Daily measurements were merged, re-clustered, allowing reliable identification of corresponding data points. The first day served as a pre-restraint day (light-dark phase) followed by three consecutive days of restraint stress. Mice were exposed to 90 min of restraint stress in a falcon tube at the end of the light phase. Due to the tetrode implantation, a longitudinal gap was made on the top of the falcon tube. Immediately after the restraint stress, sleep architecture was measured during the first 6 h in the dark phase.

At the end of the experiments, animals were transcardially perfused, and coronal sections were cut from the paraformaldehyde-fixed brains. The position of the optrodes and EEG screw were verified with help of the electrode track.

### Acute slice electrophysiology

For acute slice preparation, mice (vGAT-Cre, 3-month-old, N_AMY_=5, N_NAc_=4) were decapitated under deep isoflurane anesthesia. The brain was removed and placed into an ice-cold cutting solution, which had been bubbled with 95% O2–5% CO2 (carbogen gas) for at least 30 min before use. The cutting solution contained the following (in mM): *205 sucrose, 2.5 KCl, 26 NaHCO_3_, 0.5 CaCl_2_, 5 MgCl_2_, 1.25 NaH_2_PO_4_, 10 glucose*, saturated with 95% O_2_–5% CO_2_. Coronal slices of 300 µm thickness were cut using a Vibratome (Leica VT1000S) and placed into an interface-type incubation chamber for recovery submerged in standard artificial cerebrospinal fluid (ACSF) containing the following (in mM): *126 NaCl, 2.5 KCl, 26 NaHCO_3_, 2 CaCl_2_, 2 MgCl_2_, 1.25 NaH_2_PO_4_, 10 glucose*, saturated with 95% O_2_– 5% CO_2_. During measurements, slices were transferred individually into a submerged-type recording chamber with a superfusion system allowing constantly bubbled (95% O_2_–5% CO_2_) ACSF to flow at a rate of 3-3.5 ml/min. The ACSF was adjusted to 300-305 mOsm. All measurements were carried out at 33 –34°C, temperature was maintained by a dual flow heater (Supertech Instruments). The pipette solution contained (in mM): 110 D-gluconic acid potassium salt, 4 NaCl, 20 HEPES, 0,1 EGTA, 10 phosphocreatine di(tris) salt, 2 ATP magnesium salt, 0.3 GTP sodium salt, pH: 7.3, 280-300 mOsm and biocytin (Sigma). Pipette resistances were 3-6 MΩ when filled with pipette solution. Visualization of slices and selection of retrogradely labelled AMY or NAc projecting PVT cells was guided by AAVrg-CAG-GFP signal (see: animals/Viral injection) and was imaged under an upright microscope (BX61WI; Olympus, Tokyo, Japan equipped with infrared-differential interference contrast optics and a UV lamp). Only cells located deeper than ~50 µm (measured from the slice surface) were targeted. All cells were recorded in voltage-clamp mode and held at −65 mV holding potential during the formation of the gigaseal. Series resistance was constantly monitored after the whole-cell configuration was established, and individual recordings taken for analysis showed stability in series resistance between a 5% margin during the whole recording. After the whole-cell configuration was established, cells were clamped at −10 mV holding potential in order to record inhibitory postsynaptic currents (IPSCs). In order to evoke IPSCs, trains of 5*5ms laser light pulses (473 nm, calibrated to ~10 mW at slice surface, Sanctity Laser) were delivered at 1, 5, 10, and 20 Hz frequency by an optic fiber positioned above the slices. Recordings were performed with a Multiclamp 700B amplifier (Molecular Devices). Data was digitized at 20 kHz with a DAQ board (National Instruments, USB-6353). All data were processed and analyzed off-line using standard built-in functions of Python 2.7.0. environment. A threshold for IPSC peak detection was calculated for each neuron from a 2-minute-long baseline before stimulation trains. The threshold value was set to be 3*SD_mean_ of the signal. IPSPs were considered to be evoked (eIPSCs) if the peak was detected within the 2-10 ms range after a given stimulus on-set. Spontaneous IPSCs were detected for quantification from the 2-minute-long baseline region before stimulation trains.

### Neural data processing

#### LFP analyses

The natural sleep and anesthetized measurements were analyzed in a similar way. The LFP/EEG/EMG signals were downsampled at 3000 Hz. The power of delta (1–4 Hz), theta (4-8 Hz), sigma/spindle activity (9–14 Hz), alpha (8–13 Hz), beta (13-30 Hz) frequency bands were differentiated^89^. For the further analysis the signals were filtered on a given frequency, thereafter were the absolute value of the filtered signals calculated, on which we performed smoothing in 100 ms window (RMS; root-means square) (**Fig. 2o**, Extended Data Fig. 1a,3f). Sleep–wake states were determined by using RMS. In the natural sleep conditions, awake state was characterized by high alpha, beta activity (alpha, beta RMS) and low delta power (delta RMS) with high EMG (threshold >0.8), while sleep was characterized by low alpha and high delta activity with low EMG, and it was further subdivided into NREM and REM state. For the further classification the EMG signal was normalized. REM was determined as a high sigma/delta (sigma/delta RMS) ratio associated, with low delta and high theta power and low EMG (threshold <0.5). NREM was determined as a low sigma/delta ratio associated with high delta power or spindle activity and low EMG (threshold <0.8). NREM sleep was not divided into further subsections. In the anaesthetized conditions, desynchronized state was characterized by high alpha, beta activity (alpha, beta RMS) and low delta power (delta RMS), while sleep was characterized by low alpha and high delta activity, and it was further subdivided into NREM-like and REM-like state. REM-like states were determined as a high sigma/delta (sigma/delta RMS) ratio associated with low delta and high theta power. NREM was determined as a low sigma/delta ratio associated with high delta power or spindle activity. All animals and measurements had a different sigma/delta RMS ratio, thus, the ratio was calculated for each trial and for each animal; the pre-stimulus ratio was compared to the measurement specific median. An 18 s time window, the baseline (sleep) period (between −20 s and −2 s), was used to determine pre-stimulus sleep stages. During spontaneous sleep architecture monitoring, the oscillation fluctuation was monitored in 5 s time windows.

The delta and theta RMS changes were calculated for the normalized signal after the optical stimulation or for the spontaneous state. The starting point of the elevated delta and theta RMS segment was determined by the time point, when the normalized signal exceeded the baseline for the first time after the stimulus, while the end point of the changes was the return of the RMS signal to baseline. The lengths of the delta and theta RMS changes (recovery-time) were given by the time period between these two points. The REM onset was given by the time when all the criteria of REM state appeared at the same time. 10s 1/10 Hz 10 mW optogenetic activations were applied to analyze changes in LFP activity (delta and theta RMS, FFT, modulation index (MI)). When the sleep was ‘light’ (the EEG was desynchronized) for longer periods, the stimulation protocol was paused. Sham stimuli were used as the control. The starting point of a sham stimulus was the half point of the ISI. Sleep–wake states were determined in the case of the sham stimuli, too.

Under natural sleep conditions, the FFT and RMS changes were calculated for the frontal EEG signal. In anesthetized measurements, we could distinguish superficial and deep layers due to the insertion path of the extracellular electrode. Based on the distribution of the PVT_→NAc_ and PVT_→AMY_ fibers, as well as the location of the intertelenchephalic and corticothalamic/pyramidal track neurons in the PFC^87^, layers 1, 2-3 and 5a were taken as the superficial, while layers 5b and 6 were considered as the deep territories. From each animal, one channel per superficial vs. deep layer was further analyzed. The channels were selected based on the best signal-to-noise ratio and were located at least 300 µm away from each other. For identifying the state-dependent firing rate for PVT cells, EMG onset as an indicator for sleep/wake transition was set as described above. Awake periods were only accepted when these were preceded by a 30 s sleeping phase and were longer than 500 ms, defined as the lower limit for minimal arousal^25^. Peri-event time histograms (PSTH) were defined for each cell around the detected EMG onsets. Z-score values of the firing rates were given upon PSTH calculation for each cell to a 20 s baseline (sleep) period (between –30 s and –10 s, calculated from the onset of the EMG signal). Significant changes of the firing rates were defined upon at least two significant (z > 1.92, P > 0.05) neighboring z-score (1 s) bins in the [–10, 10]-s interval around EMG onset. Mean z-scores for PVT_→NAc_ or PVT_→AMY_ neurons are presented^25^. To determine the sleep state specific EMG changes, distinct calculation methods were used. Each animal and measurement had different EMG levels, thus for the EMG onset latency and duration were calculated for every trial, and for each animal; the pre-stimulus ratio was compared to the measurement specific median. If the median pre-stimulus EMG level was lower than the post-stimulus EMG level for each stimulus up to 18 s baseline (sleep) period (between –20 s and –2 s), then it was determined as an EMG onset.

To validate the probability of stimulation-induced state transition, we analyzed the RMS changes. The probability was assessed when the animal transitioned from one state (X) to other (Y) within a fixed time window (t = 10 s). First, all stimulation events (p) were identified during which the animal was in state X, then the number of events (q) were determined in which the animal entered state Y. State Y was considered valid if it was maintained for the entire time window or exceeded it (d). The cumulative probability of transitions were calculated according to the following equation (P(X → Y | t ≤ d) = q / p), in order to quantify the likelihood of stimulation-induced state transition (**Fig. 2o**). To validate the efficiency of stimulation-induced theta modulation, an efficiency index was calculated. This index was defined as the product of the probability of transition (see above) and the duration of achived state (in secundum): efficiency index = transition probability * state duration (**Fig. 2o**).

#### Single cell analyses

The natural sleep and anesthetized measurements were analyzed in a similar way. Noise filtering by average subtraction and raw electrophysiological recordings were filtered (>500 Hz) for spike detection. Spike detection and principal-component-analysis-based automatic clustering were performed using SpikeDetekt and KlustaKwik (2019 edition) respectively^90^. Cell grouping was manually refined using KlustaViewa (2019 edition). A group of spikes was considered to be generated by a single neuron if they formed a discrete, well-isolated cluster and had an autocorrelogram with a refractory period (if average bin values of the first 1.5 ms did not reach the autocorrelogram’s asymptote line). Cells were merged in those cases where units had the same autocorrelogram and a similarly shaped action potential on the same channel points to avoid enumerating the same cell more than once. Further data analyses were carried out using custom-built Matlab routines. All the presented electrophysiological data was derived from individually identified single units (clustered cells).

Putative principal neurons (glutamatergic cortical neurons) and GABAergic cortical interneurons were separated by the spike half-width, the autocorrelogram^91^ and the baseline firing rate ^92^, which we considered as the most reliable parameters for this purpose. Based on previous data on cortical GABAergic interneurons, we distinguished fast spiking (FS) and non-fast spiking (NFS) interneurons based on their spike half-width, baseline firing rate and autocorrelogram (Extended Data Fig. 10a-d).

We calculated the signal-to-noise ratio to determine the strength of the evoked feed-back inhibition on the cortical signal (**Fig. 5e,h**). To define it, we calculated the variance of the evoked response between two time points: T_0_ and T_evoked MUA peak_. The T_evoked MUA peak_ is the positive peak of the MUA for the 1s 10 Hz optical stimulation. The signal-to-noise ratio was then estimated as the mean at T_0_ and T_evoked MUA peak_ divided by standard deviation at T_0_ and T_evoked MUA peak_. The evoked signal-to-noise ratio was calculated for the first three evoked MUA peaks, which were compared to the signal-to-noise ratio of the stage before the stimulus (−100 ms-0 ms).

Significant changes in evoked responses of PVT cells were validated with MI. MI was calculated for the firing rate (FR) from the mean PSTH of the pre period (between –10 s and 0 s) which was compared to the mean PSTH of the post period (between 0 s and +10 s). We expressed the modulation index as the ratio of firing rate: (FR_post_ – FR_pre_)/(FR_post_ + FR_pre_)^93^. Neurons with MI between 0.2 and 1 were considered as activated and with MI between −0.2 and −1, as inhibited. Both the activated and inhibited neurons were assigned to the modulated PVT (PVT_MOD_) population. Based on these, neurons with MI between 0.2 and −0.2 were assumed as non-modulated cells (PVT_NON_; **Fig. 1i,j**).

Furthermore, in the chronic and anaesthetized data, thalamic unit activity was correlated with theta waves (phase-locking analysis in **Fig. 1l and 2k**). Based on the dominance of the distinct frequency band (delta, theta, sigma, gamma, EMG) the sleep stages were automatically scored. The different oscillations were normalized to the maximum power. REM was assigned to suprathreshold theta and subthreshold delta power, with low EMG. The sorted REM stages are cut out from the full measurement. After low-pass filtering the peaks of theta waves were defined. We have defined theta waves within a 0.18 s width value and with maximum 100-microvolt amplitude. Based on the width of waves the start and the end point of the theta waves are calculated. Using this data, sinusoidal waves were fitted and these hypothetical sinusoidal waves were phase-locked to the spiking activity of thalamic neurons.

### Statistical analysis

The number of analyzed slices and number of cells are shown with N, while n represents the number of mice. Data from independent experiments or animals were pooled when it was possible. Statistical significance was assessed using Kruskal-Wallis ANOVA with Mann– Whitney U-tests, Friedman ANOVA with Wilcoxon Matched Pairs Test, Paired t-test or Student’s t-test with normality test, as well as analysis of variance (one-way ANOVA, repeated measures ANOVA) to analyze the axon density data or the electrophysiological data. Normality was tested using the Shapiro-Wilk test. Pearson’s χ2 tests were used to compare the distribution of the responsive cortical cells. Post hoc test is used for ANOVA analysis (Fisher LSD or Bonferroni Correction). For correlation analysis Kendall’s τ coefficient was used to measure the ordinal association between two measured quantities. The Kolmogorov–Smirnov test was used to validate the distribution of a sample and to compare the distributions of two independent sample groups. An Equivalence Independent Samples T-test (TOST) was performed to assess the correspondence between data obtained under anesthesia and in freely moving conditions ^94,95^. Statistical analysis was performed using JASP, MATLAB or Microsoft Office Excel. Significance levels are indicated as follows: 0.1 > #P > 0.05; *P < 0.05; **P < 0.01; ***P < 0.001; ****P < 0.0001. No statistical methods were used to predetermine sample size, but it is comparable to previously published work^25,96,97^.

## Acknowledgements

We thank Z.J. Huang and László Acsády for providing us transgenic mice, Tamás Herczeg, Réka Erdős, Anna Fehér, and Katalin Varga for laboratory assistance, Csaba Dávid and Ákos Babiczky for advising us on axonal analysis and Gábor Nyíri for sharing primary antibodies. We thank Tallie Z. Baram, Antoine Adamantidis, Nadia Urbain and Iván Soltész for comments and discussions about the manuscript. We also thank the Institute of Molecular Life Sciences of the HUN-REN Research Centre for Natural Sciences for the use of Zeiss microscope and we would like to thank the Light Microscopy Center of HUN-REN Institute of Experimental Medicine. We also thank the Virus Technology Unit of HUN-REN Institute of Experimental Medicine. This work was supported by the following funding agencies: Ministry of Innovation and Technology of Hungary from the National Research, Development and Innovation Fund (K138836, and KKP126998 to FM; FK135285, FK144583 to SB; PD124034 to BB; HU-RIZONT-2024-00003 to AM); Hungarian Brain Research Program (2017-1.2.1-NKP-2017-00002 to FM; NAP2022-I-7/2022 to MLL); New National Excellence Program of the Ministry for Innovation and Technology (ÚNKP-21-5-ÁTE-2 to FM; ÚNKP-22-2-III-ELTE-539 to AVB, ELKH SA-48/2021); National Natural Science Foundation of China (8234101126 to DL); Cooperative Doctoral Program (KDP-2020-1015461 to AM); New national excellence program of the ministry for culture and innovation from the source of the national research, development and innovation fund (EKÖP-2024-87 to PB, EKÖP-2024-276 to AVB, 2025-2.1.1-EKÖP-2025-00014 to AM). FM was a János Bolyai Research Fellow.

## Contributions

A.M., P.B., B.B. and F.M. conceived and designed the project. A.M., P.B., B.B., A.V.B. and F.M. performed experiments. A.M., P.B., S.B. and F.M. analysed data. S.B., M.L.L and F.M. supervised electrophysiology experiments. F.M. supervised the entire project. A.M., P.B., M.L.L., D.L. and F.M. prepared figures and wrote the original draft. The other authors made valuable comments on the manuscript and figures.

## Data availability

All data that support the findings of this study are available from the corresponding author upon request. Source data are provided with this paper.

## Code availability

Custom-written codes used to analyze data from this study are available from the corresponding author upon request.

