## supplementary material for "A thalamic hub integrates brainstem and stress signals to dynamically regulate REM sleep"

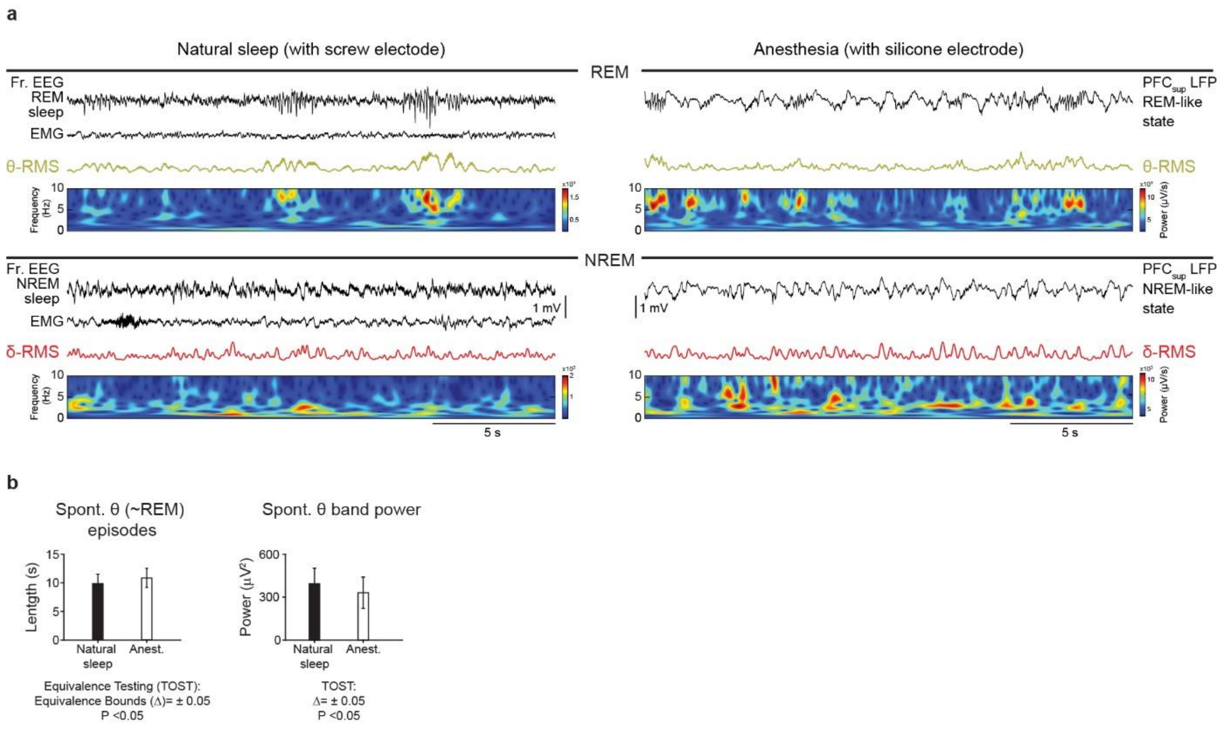


**Extended Data Fig. 1: Drug-free (natural) and anesthetized experimental design used in this study provided comparable sleep oscillation characteristics**

**a,** Comparable slow oscillation profiles during drug-free (natural) sleep and urethane anesthesia. Note similar RMS profiles for REM and REM-like as well as for NREM and NREM like states, respectively. **b,** Similarity in duration of spontaneous REM and REM-like periods with high theta oscillation (*left*) and theta band power (*right*) during natural sleep and anesthesia analyzed with Equivalence Testing (Two One-Sided Tests, TOST). (Also see ***Methods***)

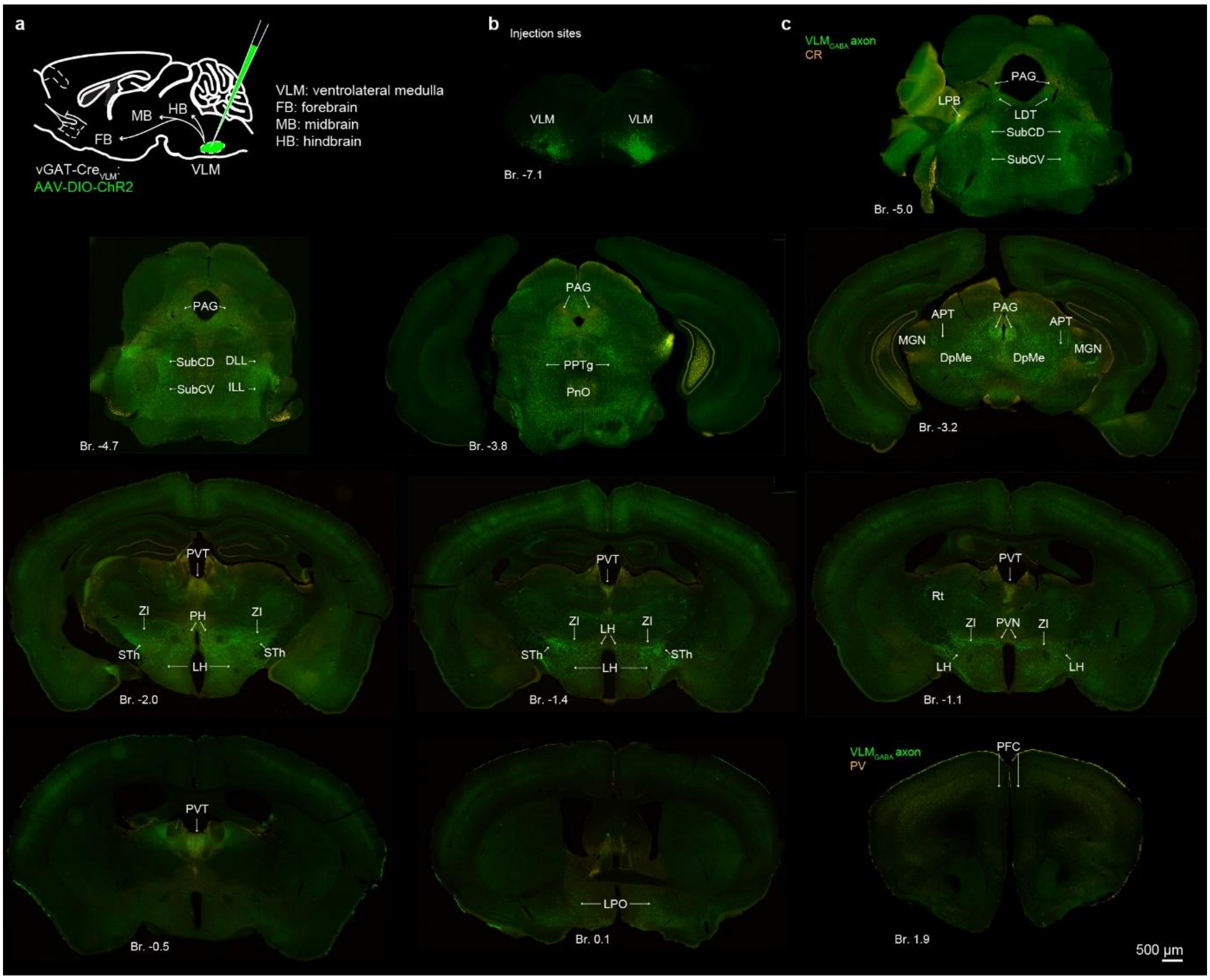


**Extended Data Fig. 2: Brain-wide axonal projections of VLM$_{GABA}$ neurons**.

**a,** Experimental design for investigating the brain-wide efferent connections of VLM$_{GABA}$ neuronal population using viral anterograde tracing in vGAT-Cre mice. **b,** Confocal image showing the injection sites of an example trial. **c,** Confocal images showing the brain wide distribution of the eYFP-labeled VLM$_{GABA}$ axonal projections.

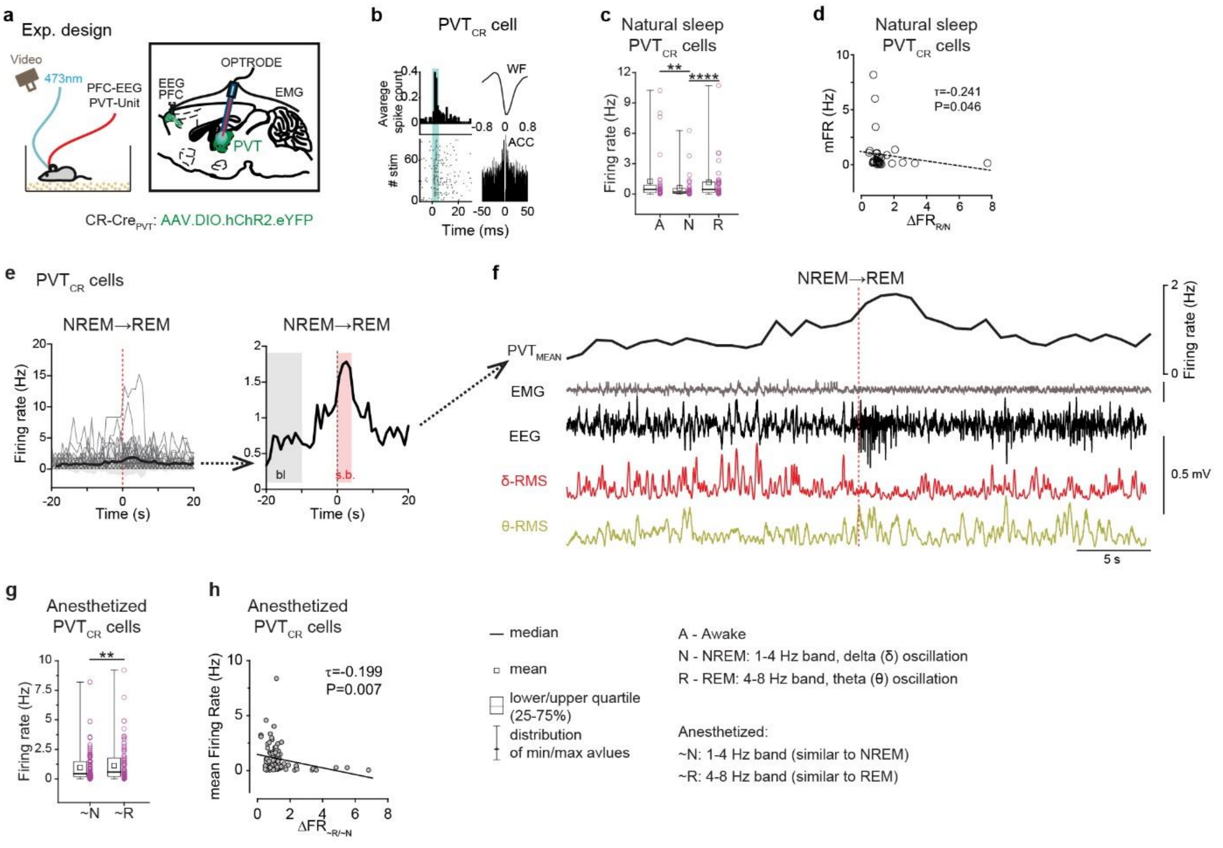


**Extended Data Fig. 3: Sleep-stage dependent activity of $PVT_{CR}$ neurons during natural sleep and anesthesia.**

**a,** Experimental design for investigating the activity of $PVT_{CR}$ cells across distinct sleep stages. **b,** Optogenetic identification (blue) of a $PVT_{CR}$ cell. WF, waveform; ACC, autocorrelogram. **c,** State-dependent firing rates of $PVT_{CR}$ cells during natural sleep condition. (N= 34 cells, n = 3 animals; Friedman ANOVA with Wilcoxon matched-pairs test). A, awake; N, NREM sleep; R, REM sleep. **d,** Negative correlation between the mean firing rate (mFR) and REM-induced firing rate changes ($\Delta FR_{R/N}$) of individual $PVT_{CR}$ cells during natural sleep. **e,** *Left*, firing rate changes of individual $PVT_{CR}$ cells around the NREM-to-REM transition during natural sleep. Red dashed line, REM onset; black line + shadow, mean ± s.d.; grey lines, activity of individual $PVT_{CR}$ cells. *Right*, averaged REM-linked increase in $PVT_{CR}$ activity prominently observed at the NREM-to-REM transition during natural sleeping (Friedman ANOVA with Wilcoxon matched-pairs test). Pink, 1 s bins with significant elevation (s.b.) compared to baseline (bl, grey). **f,** An example 40 s-long (-20 s – to 20 s) trace of EMG and EEG with corresponding delta and theta RMS signals showing a NREM→REM switch (red dashed line) along with the mean firing rate of the $PVT_{CR}$ population (thick black line; same as in E). **g,** State-dependent firing rates of $PVT_{CR}$ cells in anesthetized condition (N = 84 identified cells, n = 7 mice). **h,** Negative correlation between mean firing rate and REM-induced firing rate changes (R/N) of individual $PVT_{CR}$ cells is also present under anesthesia, similar to natural sleep condition. $^{**}P < 0.01$; $^{****}P < 0.0001$. (Statistics in **Extended Data Table 1**).

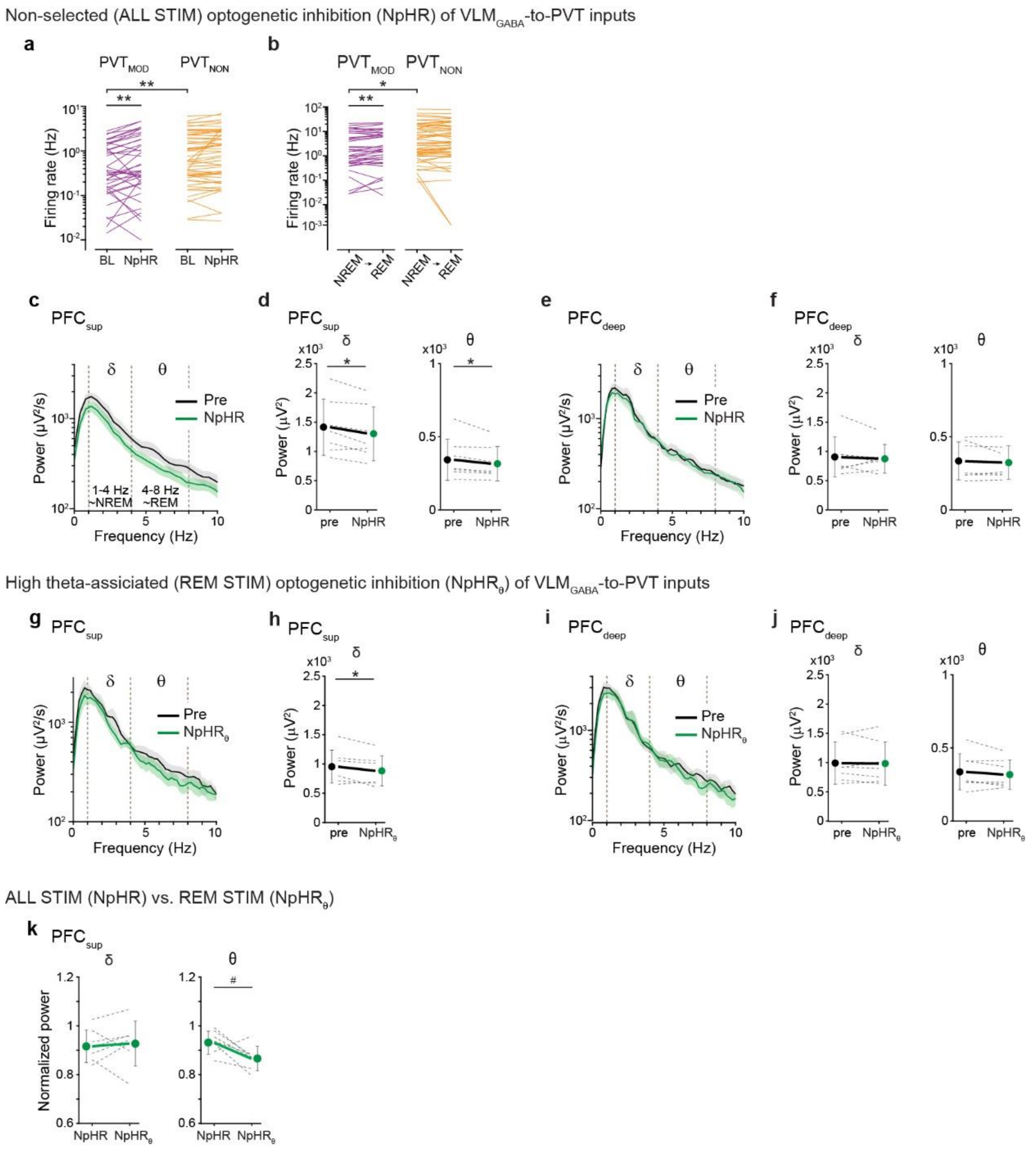


**Extended Data Fig. 4: Further analyses of VLM$_{GABA}$ influence on PVT cellular and PFC LFP activity (related to Fig. 1).**

**a,** Baseline (BL) and evoked activity of PVT cells followig inhibition of VLM$_{GABA}$-to-PVT inputs in ALL STIM trials (NpHR), clustered by responsiveness to the inhibition [PVT$_{MOD}$ (N = 40 cells) vs. PVT$_{NON}$ (N=55 cells)]. **b,** Firing rate changes upon NREM-to-REM transition were observed only in the PVT$_{MOD}$ group (in ALL STIM trials). **c-j**, FFT changes and quantification of LFP responses to inhibition of VLM$_{GABA}$-to-PVT inputs (green) in PFC$_{sup}$ and PFC$_{deep}$, compared to pre-stim (black) in *ALL STIM* **c-f**, and in *REM* STIM trials (**g-j**); (related to Fig. 1l). **k,** Comparison of LFP responses in ALL STIM vs. REM STIM trials. $^{\#}P < 0.1$; $^{*}P < 0.05$; $^{**}P < 0.01$. (Statistics in **Extended Data Table 1**).

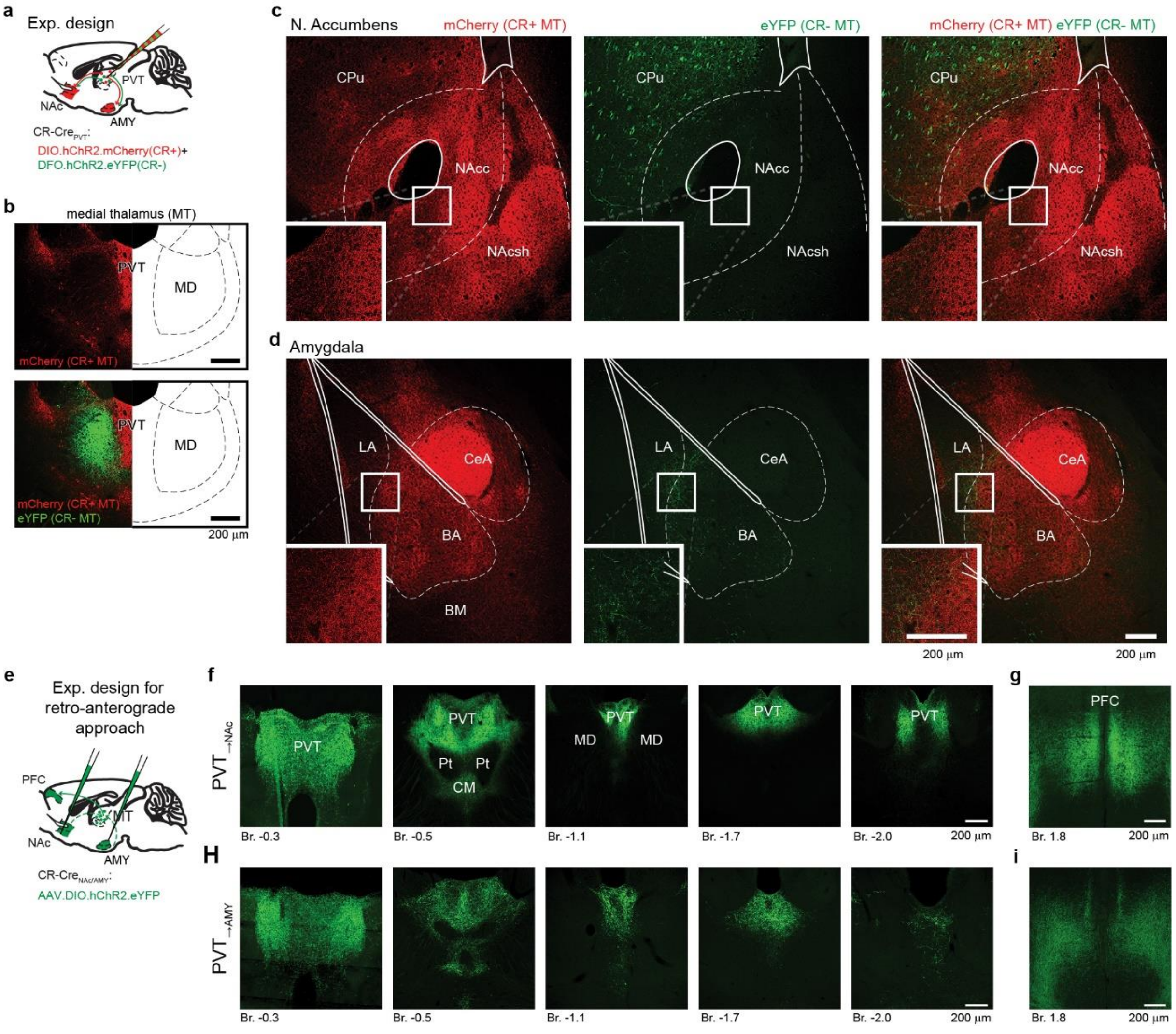


**Extended Data Fig. 5: Anatomical validation of CR-Cre and retro-anterograde labeling for targeting $PVT_{\rightarrow NAc}$ and $PVT_{\rightarrow AMY}$ neurons (related to Fig. 2).**

**a,** Experimental design for selective viral labeling of CR+ (PVT; red) and CR- medial thalamic (MT) cells (mediodorsal thalamic; MD; green). **b,** Confocal images of the injection site in MT. **c,d,** Confocal images showing strong CR+ (mCherry, red) but weak or no CR- (YFP, green) axonal innervation in the nucleus accumbens (**c**) and the amygdala (**d**). Insets represent the highest density of CR+ and CR- thalamic fibers at higher magnification. **e,** Experimental design for retroanterograde labeling of $PVT_{\rightarrow NAc}$ and $PVT_{\rightarrow AMY}$ neurons. **f,h,** Confocal images representing the distribution of retrogradely labeled $PVT_{\rightarrow NAc}$ (YFP; **f**) and $PVT_{\rightarrow AMY}$ neurons (YFP; **h**) along the entire anteroposterior extent of PVT. **g,i,** Confocal images representing cortical collateral axons of retrogradely labeled $PVT_{\rightarrow NAc}$ (YFP; **g**) and $PVT_{\rightarrow AMY}$ neurons (YFP; **i**) in PFC using the retro-anterograde labeling methods (see also ***Methods***).

Abbreviations: BA, basolateral amygdala; BM, basomedial amygdala; CeA, central amygdala; CM, centromedial thalamus (th); CPu, caudate putamen; LA, lateral amygdala; MD, mediodorsal th; NAcc and NAcsh nucleus accumbens core and shell; Pt, paratenial th.; PVT, paraventricular th..

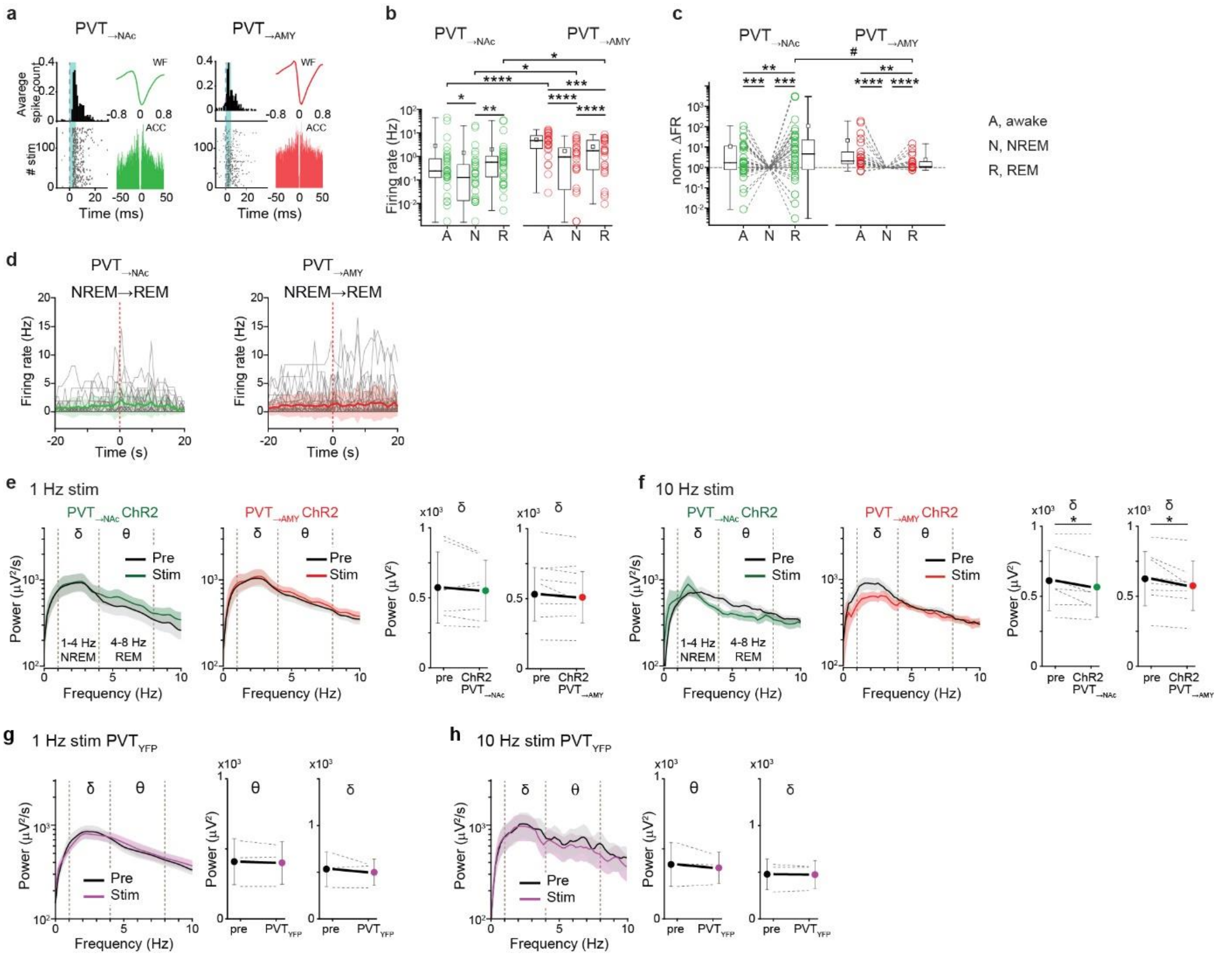


**Extended Data Fig. 6: Further analyses of PVT→NAc neuron-mediated VLMGABA effects on REM sleep (related to Fig. 2).**

**a,** Optogenetic tagging of PVT→NAc and PVT→AMY. F, waveform; ACC, autocorrelogram. **b,c,** Sleep state-dependent firing rates of PVT→NAc (N = 25 cells, n = 7 mice) and PVT→AMY (N = 36 cells, n = 8 mice) cells during natural sleep shown with original (**b**) and normalized (**c**) values (Wilcoxon matched-pairs test for comparing states-dependent firing rates; Mann-Whitney U test for PVT→NAc vs. PVT→AMY). **d,** Firing rate changes of individual PVT→NAc and PVT→AMY cells around the NREM-to-REM transition during natural sleep. Red dashed line, REM onset; green/red lines + shadows, mean ± s.d.; grey lines, individual neuronal data. **e,f,** *Right*, frequency-dependent FFT changes evoked by 1 Hz (**e**) and 10 Hz (**f**) optogenetic stimulation of PVT→NAc and PVT→AMY neurons. *Left,* population data for delta (δ) band power changes induced by PVT→NAc and PVT→AMY stimulation (n = 7-8 animals; Wilcoxon matched-pairs test). **g,h,** 1 Hz (**g**) and 10 Hz (**h**) optogentic activation of control, YFP-transduced PVT neurons (n = 3 mice) did not elicit changes in sleep oscillations. #P < 0.1; *P < 0.05; **P < 0.01; ***P < 0.001; ****P < 0.0001. (Statistics in **Extended Data Table 1**).

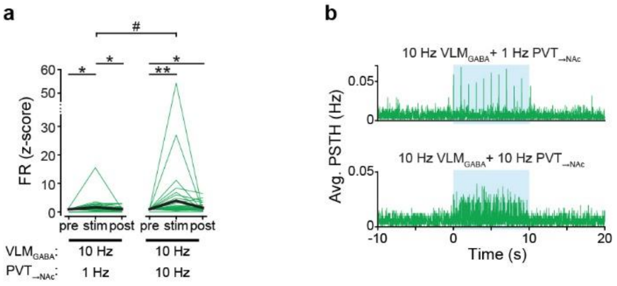


**Extended Data Fig. 7: Further analyses of the impact of $PVT_{\rightarrow NAc}$ and $VLM_{GABA}$ co-ativation on $PVT_{\rightarrow NAc}$ activity (related to Fig. 3).**

**a,** Firing rate changes of $PVT_{\rightarrow NAc}$ neurons (N=41) at 1 Hz and 10 Hz antidromic optogenetic stimulation of $PVT_{\rightarrow NAc}$ along with 10 Hz orthodromic optogenetic stimulation of $VLM_{GABA}$ neurons (Wilcoxon matched-pairs test), presented as z-scores (Wilcoxon matched-pairs test for comparing firing rates during pre, stim and post periods; Mann-Whitney U test for comparing 1 Hz vs. 10 Hz). **b,** Peristimulus time histograms (PSTHs) showing averaged activity changes of $PVT_{\rightarrow NAc}$ neurons evoked by $VLM_{GABA}$ and $PVT_{\rightarrow NAc}$ optogenetic co-activation. $^{\#}P < 0.1$; $^{*}P < 0.05$; $^{**}P < 0.01$. (Statistics in **Extended Data Table 1**).

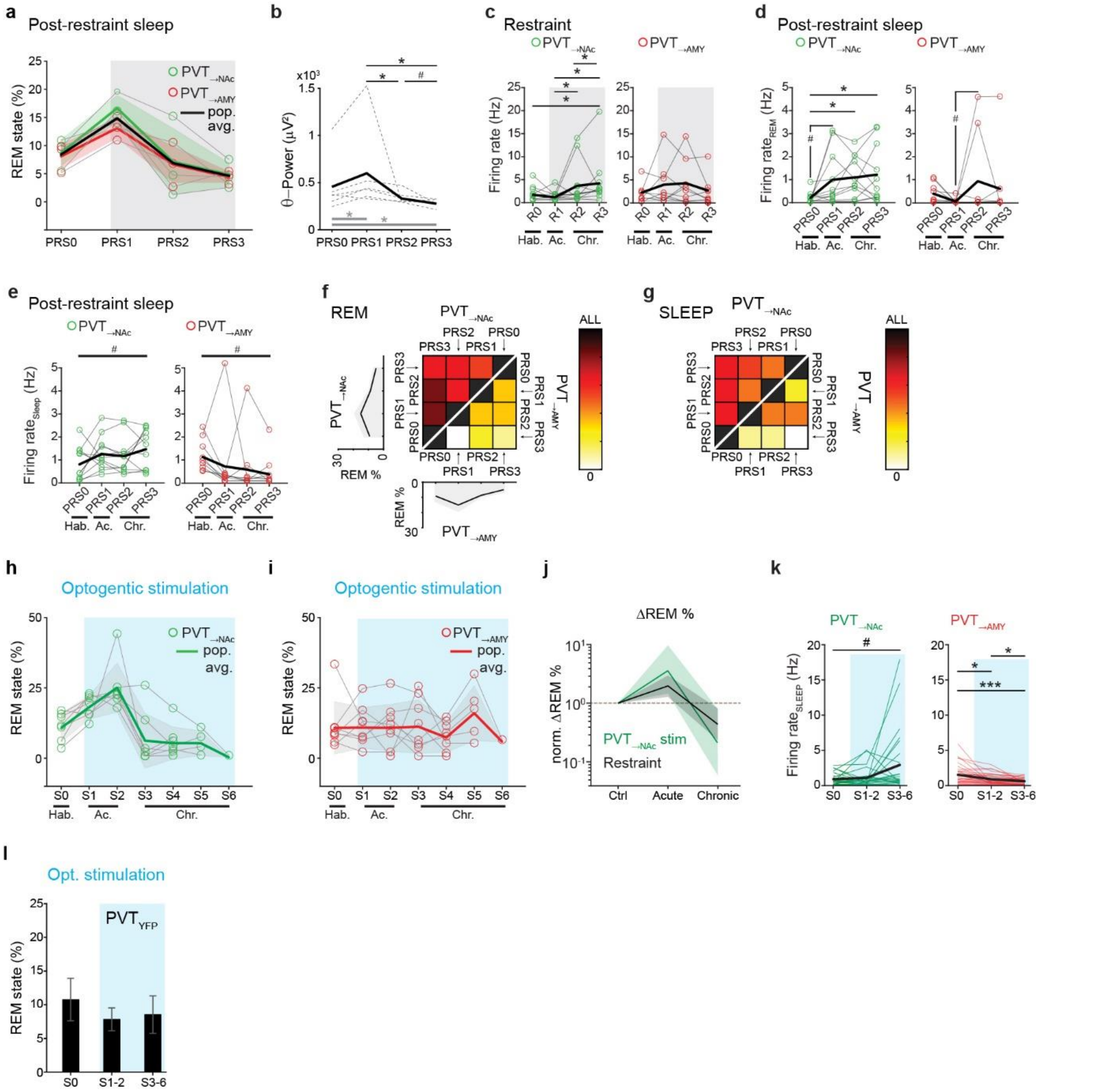


**Extended Data Fig. 8: Further data on stress-induced bidirectional REM modulation by PVT$_{\rightarrow NAc}$ neurons (related to Fig. 4).**

**a,** Restraint-induced REM sleep changes of individual animals (PVT$_{\rightarrow NAc}$ n = 3; PVT$_{\rightarrow AMY}$ n = 3) during post-restraint periods (PRS1-3) compared to the habituation day (Hab.; PRS0). **b,** Population data for theta band changes associated with restraint-induced REM sleep alterations across experimental days (n = 6 mice; Wilcoxon matched-pairs test). **c-e,** Population data for restraint-induced activity changes of PVT$_{\rightarrow NAc}$ (*left*; N = 11 cells) and PVT$_{\rightarrow AMY}$ cells (*right*; N = 9 cells) during restraint (**c;** R0-R4), post-restraint REM sleep (**d;** PRS0-PRS3) and post-restraint total sleep (**e**). R0 and PRS0, habituation days; R1 and PRS1, acute restraint phase; R2-3 and PRS2-3, chronic restraint phase. **f,g,** Heat maps indicating the number of thalamic cells showing restraint-induced elevated activity during REM sleep (**f**) and during the entire sleep periods (**g**) compared to previous days. **h,i,** Data from individual animals during optogenetic stimulation (blue) of PVT$_{\rightarrow NAc}$ (**h**) and PVT$_{\rightarrow AMY}$ (**i**). **j,** Normalization reveals that PVT$_{\rightarrow NAc}$ activation induced REM sleep changes similar to those caused by acute and chronic restraint stress. **k,** Population data indicate sleep-related activity changes of $PVT_{\rightarrow NAc}$ (N = 25) and $PVT_{\rightarrow AMY}$ (N= 36) neurons during non-stim periods (Repeated Measures ANOVA with post hoc LSD test). **l,** Optogenetic stimulation (blue) of YFP-transduced PVT neurons (n = 3) did not evoke REM state changes (Friedman ANOVA). $^{\#}P < 0.1$; $^{*}P < 0.05$; $^{***}P < 0.001$. (Statistics in **Extended Data Table 1**).

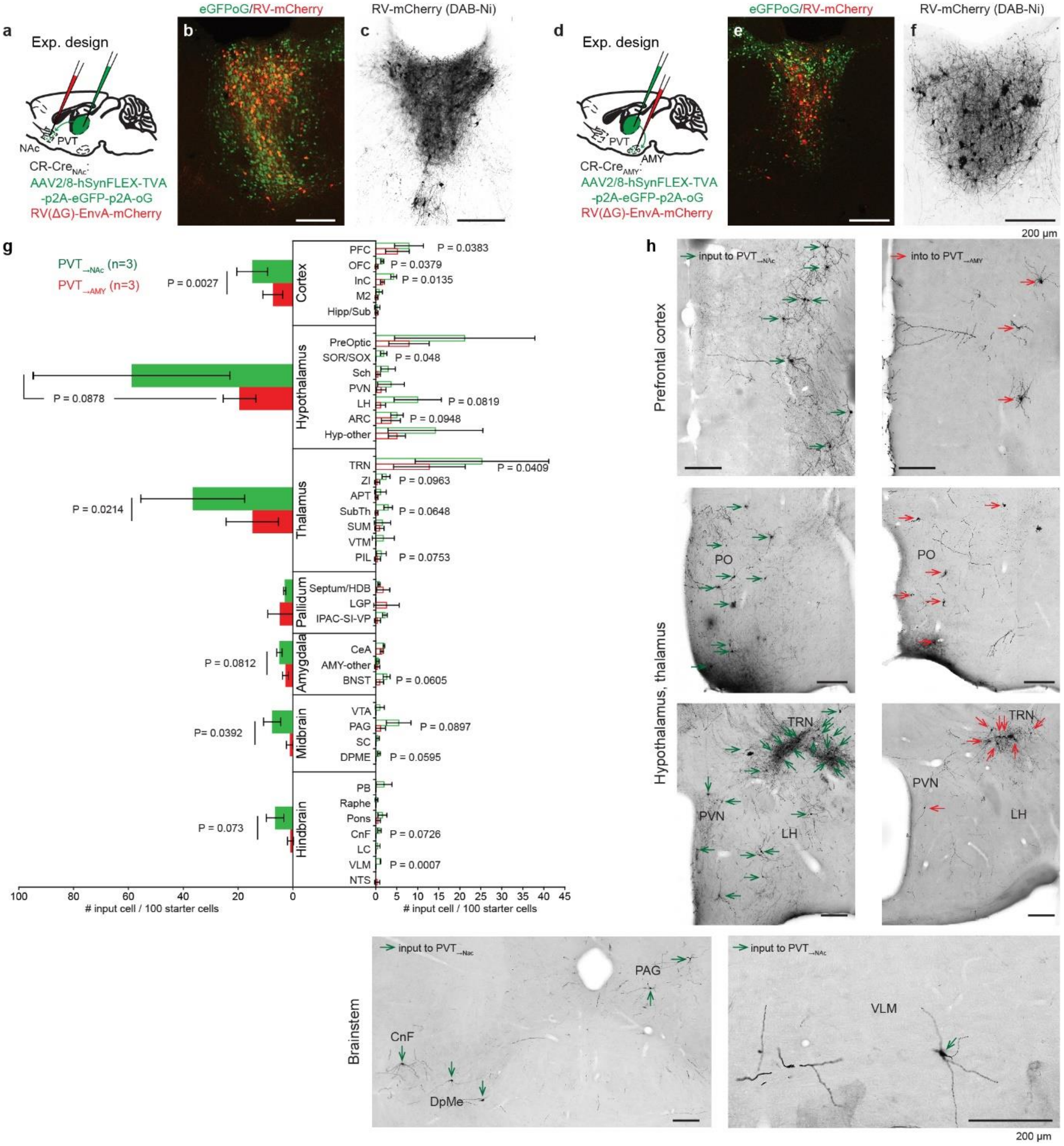


**Extended Data Fig. 9: Different brain-wide inputs of PVT→NAc and PVT→AMY cells.**
**a,d,** Experimental design for investigating brain-wide afferentation of PVT→NAc (**a**) and PVT→AMY cells (**d**) using rabies-mediated monosynaptic retrograde tracing. **b,e,** Expression of AAV2/8-hSynFLEX-TVA-p2A-eGFP-p2A-oG (green) and rabies-mediated mCherry (red) in a projection-specific manner: PVT→NAc (**b**) and PVT→AMY (**e**). **c,f,** DAB-Ni-labeled PVT→NAc (**c**) and PVT→AMY (**f**) starter cells. **g,** Population data showing the distribution of input cells of PVT→NAc (green) and PVT→AMY neurons (red). **h,** Representative micrographs showing DAB-Ni-labeled input cells of PVT→NAc (green arrows; *left*) and PVT→AMY neurons (red arrows; *right*). (Statistics in **Extended Data Table 1**).

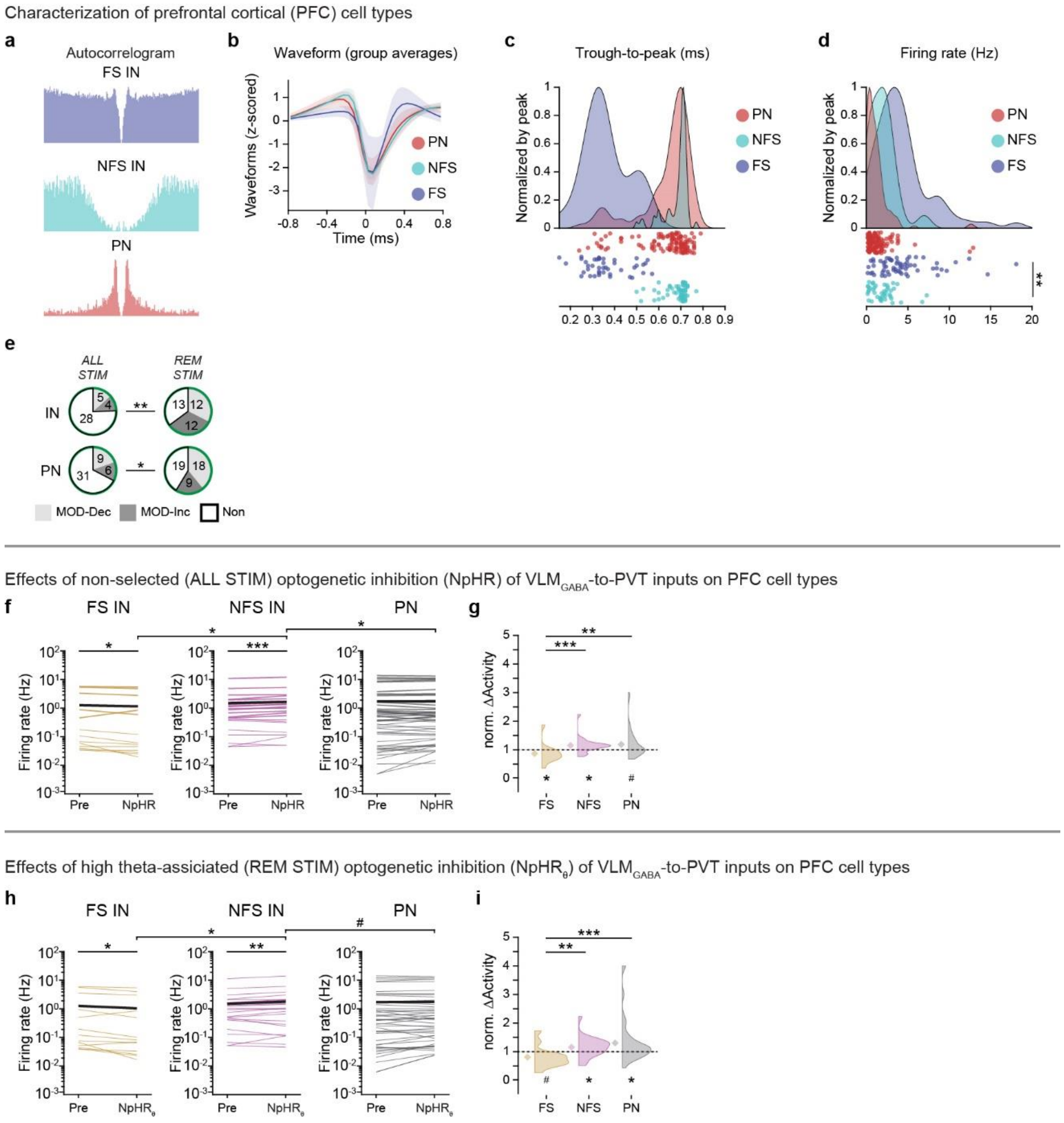


**Extended Data Fig. 10: Impacts of VLM$_{GABA}$ on PFC circuits via the PVT (related to Fig. 5).**

**a-d,** Electrophysiological characterization of PFC cell types: fast spiking interneurons (FS IN); non-fast spiking interneurons (NFS IN) and principal neurons (PN). **e,** Pie charts showing state-dependent responsiveness of PFC IN (N = 37 cells) and PN cells (N = 46 cells) to the inhibition of VLM$_{GABA}$-to-PVT inputs (n = 7 animals; Pearson's $\chi 2$ test). **f-i,** Activity changes of FS, NFS and PN PFC cells evoked by the inhibition of VLM$_{GABA}$-to-PVT inputs during *ALL STIM* (**f**,**g**) and *REM STIM* trials (**h**,**i**). **f**,**h**, original data; **g**,**i**, normalized data. (Wilcoxon matched-pairs test for Pre-stim vs. NpHR$_{REM}$; Mann-Whitney U test for FS vs. NFS vs. PN). $^{\#}P < 0.1$; $^{*}P < 0.05$; $^{**}P < 0.01$; $^{***}P < 0.001$. (Statistics in **Extended Data Table 1**).

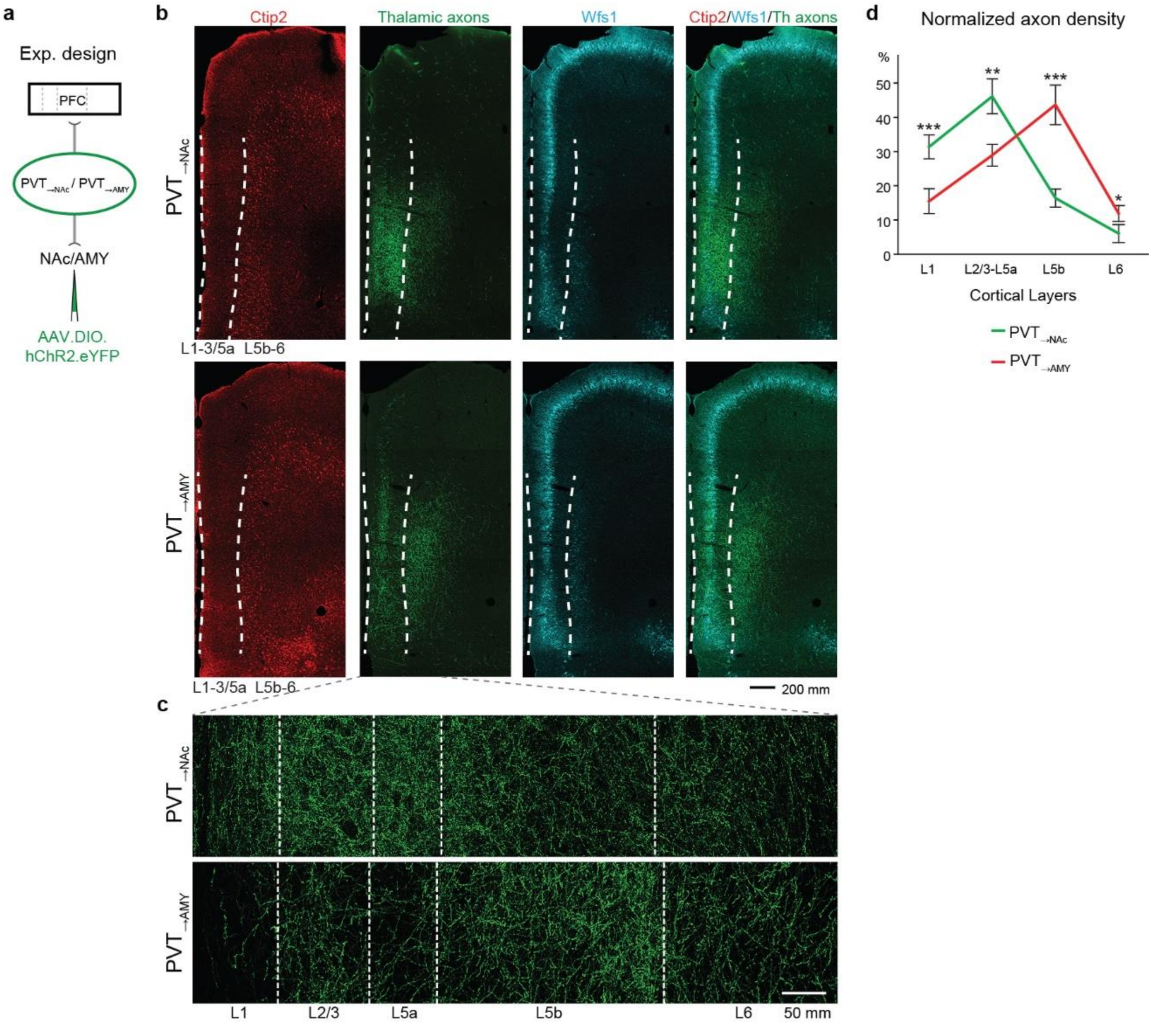


**Extended Data Fig. 11: Further data on layer selective cortical innervation by PVT→NAc and PVT→AMY neurons (related to Fig. 5).**

**a,** Experimental design for investigating the axonal arbor of PVT→NAc and PVT→AMY neurons in PFC. **b,** Layer-selective neurochemical marker labeling in the PFC overlaid with cortical innervation of PVT→NAc or PVT→AMY neurons (green). Ctip2 for layer 5b-layer 6 (red) and Wfs1 for layer 2. White dashed line represents the borders of L1-L5a (containing intratelenchephalic neurons; superficial layers) and L5b-L6 (containing corticothalamic and pyramidal tract neurons; deep layers). **c,** Representative confocal images showing the laminar distribution of PVT→NAc (*top*) or PVT→AMY (*bottom*) inputs in the PFC. **d,** Normalized axon density. *P < 0.05; **P < 0.01; ***P < 0.001. (Statistics in **Extended Data Table 1**).

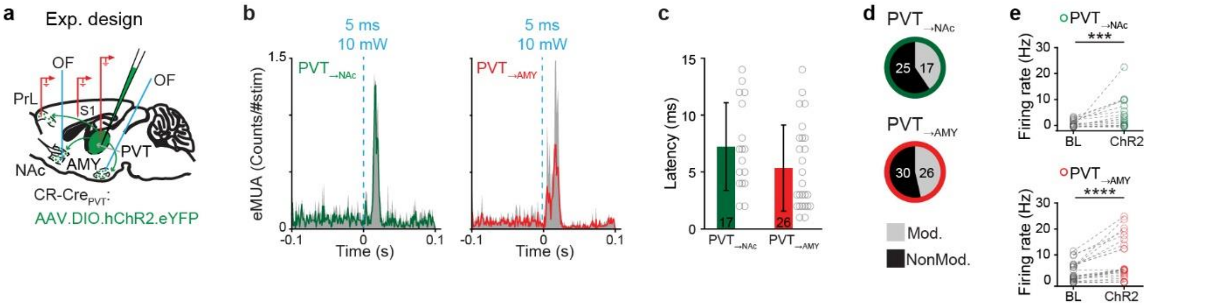


**Extended Data Fig. 12: Reliable retrograde activation of PVT→NAc and PVT→AMY neurons (related to Fig. 5).**

**a,** Schematic of in vivo recordings and retrograde stimulation of PVT→NAc and PVT→AMY cells under anesthesia. **b-e,** Reliable antidromic optogenetic activation of photo-identified PVT→NAc (N= 17 cells) and PVT→AMY (N= 26 cells) (**b**) with similar response latency (**c**), and comparable activation efficacy (**d,e**). ***$P < 0.001$; ****$P < 0.0001$. (Statistics in **Extended Data Table 1**).

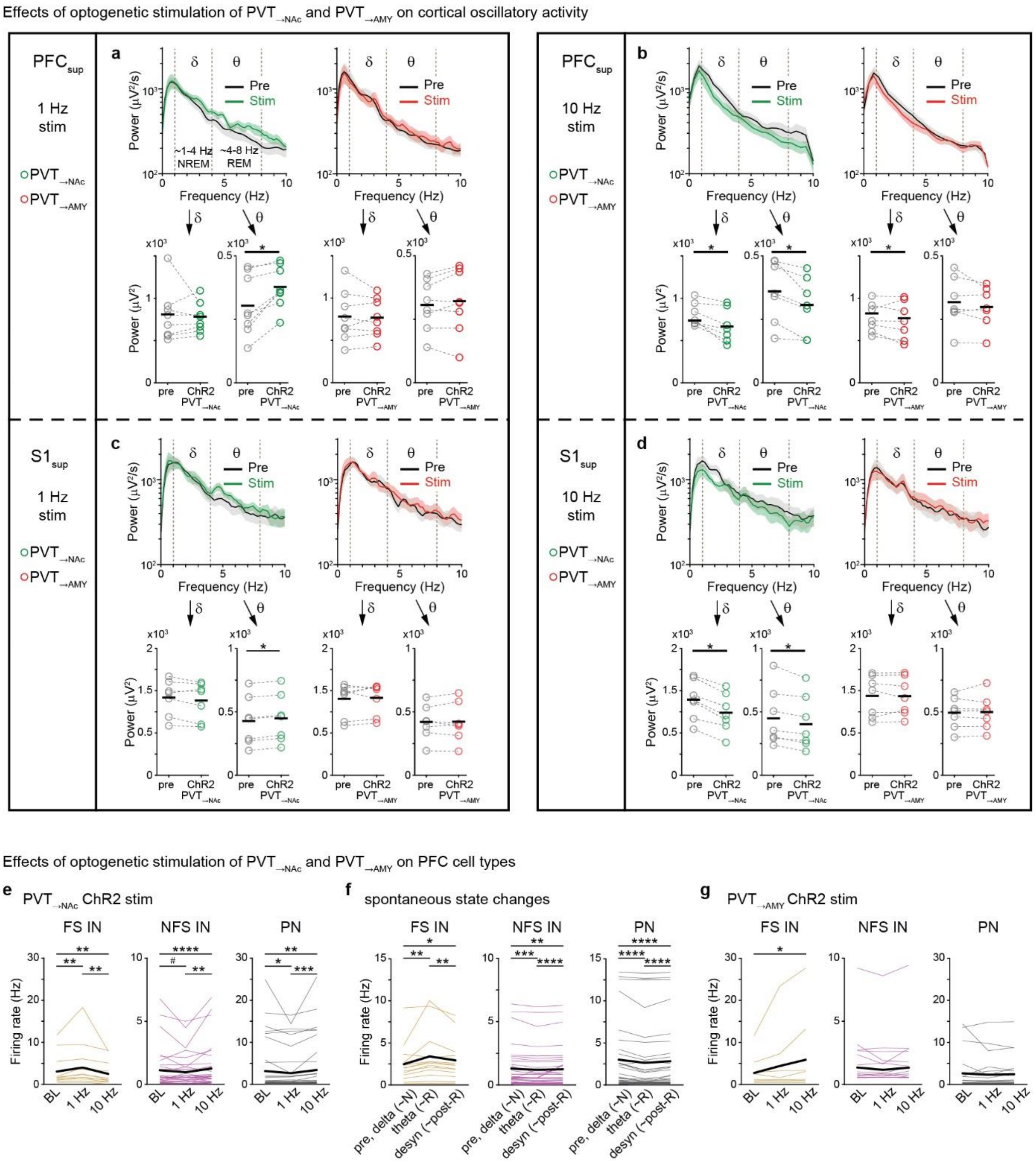


**Extended Data Fig. 13: Frequency-dependent effects of PVT→NAc and PVT→AMY activation on cortical oscillatory and single cell activity under anesthesia (related to Fig. 5).**

**a-d,** Frequency-dependent bidirectional effects of optogenetic stimulation of PVT→NAc and PVT→AMY neurons on delta and theta oscillations in the superficial layers of PFC (**a,b**) and primer somatosensory cortex (S1; **c,d**) under anesthesia. *Top*, FFT between 0-10 Hz; *bottom*, changes of FFT power of delta and theta band oscillations (Wilcoxon matched-pairs test). **e,** Opposite activity changes in PFC FS (N=13 cells), NFS (N=33 cells) and PN (N= 37 cells) populations triggered by 1 Hz and 10 Hz PVT→NAc activation; presented with original values (Wilcoxon matched-pairs test). **f,** Comparable changes in PFC FS, NFS and PN activation during spontaneous state transitions and during 1 or 10 Hz stimulation (**e**): activity changes at the shift from high-delta to high-theta periods (similar to natural NREM → REM) resemble the impacts of 1 Hz stimulation; whereas transitions from high-theta to desynchronized state (similar to natural REM → post-REM) resemble the impacts of 10 Hz stimulation (shown with original values). **g,** $PVT_{\rightarrow AMY}$ stimulation elicited different or no state-dependent activation of PFC FS, NFS and PN cells indicated with original values. $^{\#}P < 0.1$; $^{*}P < 0.05$; $^{**}P < 0.01$; $^{***}P < 0.001$; $^{****}P < 0.0001$. (Statistics in **Extended Data Table 1**).

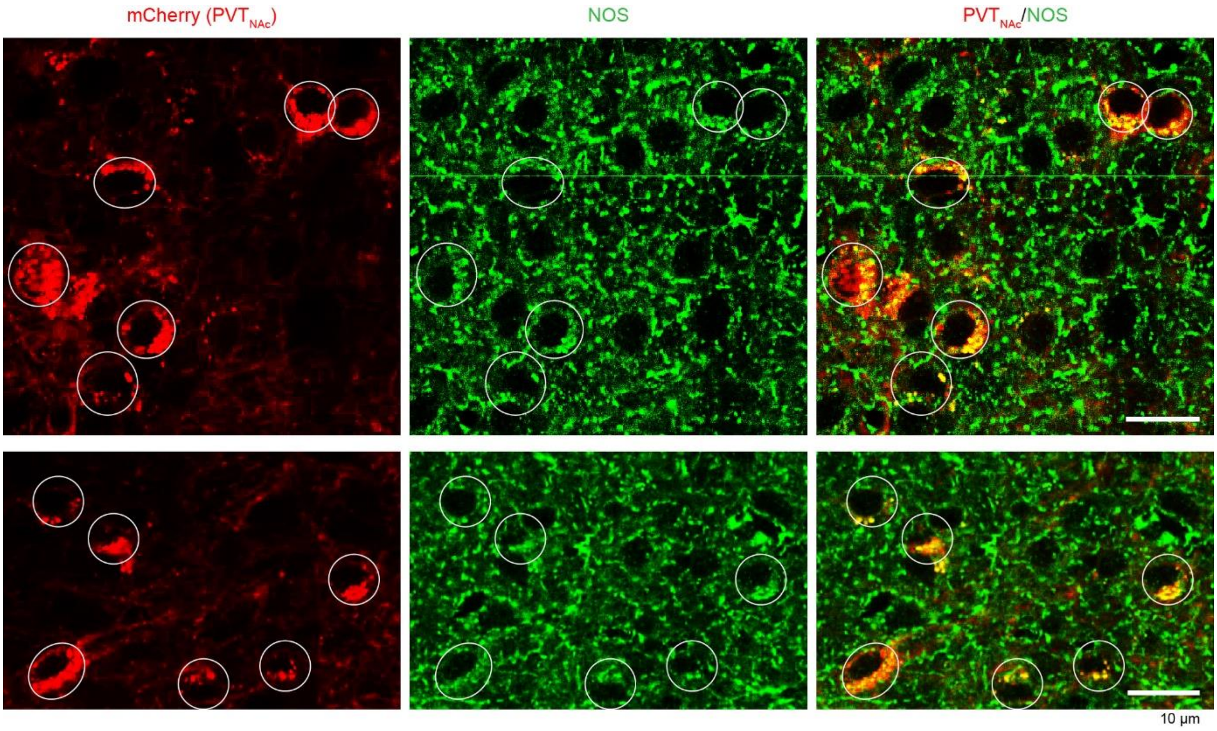


**Extended Data Fig. 14: NOS expression by $PVT_{\rightarrow NAc}$ cells.**

Two representative confocal images indicate that the retrogradely labeled $PVT_{\rightarrow NAc}$ cells by AAV-DIO-mCherry (red; *left*) express nitric oxide synthase (NOS; green; *middle*). *Right*, overlay of mCherry and NOS expressions.
